# Finding Salient Multi-Omic Interactomes Relevant to Multiple Biomedical Outcomes using Graph Ensemble Neural Networks

**DOI:** 10.1101/2024.12.28.630633

**Authors:** Tara Eicher, Jany Chan, Aline P. Becker, S. Jaharul Haque, Jessica Fleming, Joseph McElroy, Wei Meng, Amy Webb, Rachel S. Kelly, Juan Celedón, Clary Clish, Robert Gerszten, Michael McGeachie, Scott T. Weiss, Jessica Lasky-Su, Arnab Chakravarti, Ewy A. Mathé, Raghu Machiraju

## Abstract

Although multi-omics integration relevant to patient outcome is typically characterized by an analyte interactome, current multi-omic integration methods either (1) model outcome without directly including associations between analytes, (2) model the interactome without directly evaluating the saliency of the model in the context of outcome, or (3) model outcome in a high-dimensional parameter space not suitable for small sample sizes (which are common in multi-omics studies). We introduce Graph Ensemble Neural Network (GENN), a methodology that learns the interactome most predictive of outcome in a low-dimensional parameter space built on complementary attributes for all possible analyte associations (*metafeatures*). We show that GENN is robust to noise in measurements using a theoretical model, outperforms the predictive performance of existing methods when evaluated on Tegafur drug response in NCI-60 cancer cell line data, and uncovers potentially novel multi-omic mechanisms driving total serum IgE levels in pediatric asthma and patient survival in glioblastomas.

## Introduction

Discovering the interactome, or significant associations between analytes (e.g., genes and compounds) relevant to clinical outcomes aids in identifying targetable functional groups like molecular pathways. Large-scale multi-omics profiling now enables tracking health outcomes and disease etiology across diverse populations. For instance, the National Heart, Lung, and Blood Institute’s TOPMed program collects asthma data to explain how gene transcription affects increased total serum Immunoglobulin E (IgE) in pediatric asthma.

In this study, 22,772 harmonized genes and 529 metabolic analytes were analyzed from 564 and 309 human subjects, respectively. **The goal is to determine the sub-interactome, or groups of associated analytes that directly impact the total serum IgE in patients**. Notably, novel associations, i.e., inferred interactions between sphingolipids (which modulate airway inflammation), serine and threonine metabolism (precursors in sphingolipid biosynthesis), immune response and IgE production (antibody response to allergens), and mitochondrial organization (whose abnormalities exacerbate inflammation) could offer a comprehensive understanding of mechanisms driving IgE levels in pediatric asthma.

Several methods evaluate saliency of analyte subgroups in low-dimensional space, but do not model the interactome as a graph, making it difficult to determine associations within the relevant analyte subgroups. For example, the Least Absolute Shrinkage and Selection Operator (LASSO) effectively obtains a subgroup of analytes predictive of outcome and “sparsifies” the number of parameters in a linear model [6]. Group LASSO is an extension in which the weights of analytes are penalized based on membership to predefined groups (e.g., biological pathways) thus reducing number of features while improving accuracy [7]. Additionally, Random Forest (RF) combines decision trees built on subsets of the original data to predict outcome; the importance of each analyte can be measured using one of several indices which are described in the **Supplementary References #2**.

There exist methods which are graph-based and represent analytes as nodes and their associations as edges. These methods evaluate the saliency of analytes subgroups and/or subgraphs building a predictive model of outcome using the interactome, thereby limiting the graph to analytes associations relevant to outcome in a complementary way. One salient example is the *Bayesian Network* (BN), which dynamically learns the interactome by (**i**) obtaining a salient joint probability graph between analytes, other features, and outcome(s) and (**ii**) learning weights associated with each edge in the probability graph, which is often an intractable problem for large graphs with thousands of analytes, typical of multi-omics contexts [8, 9]. Proposed solutions which constrain the in-degree of each node in the graph or the selection of parent nodes for each node in the graph are still not scalable (see **Supplementary References #3**).

Another recent graph-based method is the Graph Neural Network (GNN), which is traditionally used to classify individual nodes in a graph but can also be used to classify entire graphs such as those generated from analyte measurements and associations (see **Supplementary References #4** for graph classification GNN models). A GNN accepts as input a pre-computed interactome and learns a complex non-linear function of the analytes or their features (e.g., gene expression, copy number variation, and methylation) to predict outcome using a framework of many nonlinear function modules (*perceptrons*) which feed into one another in a series of layers [10]. Notably, identification of the salient subgraph is not always straightforward. Liu et al describe several approaches [11] (**Supplementary References #5)**. Nevertheless, as the number of analytes in multi-omics studies is typically greater than the number of samples, weights learned using either BN or GNN may be unreliable due to the *curse of dimensionality problem* [12], leading to low generalizability (i.e., poor predictive performance on new data not included in the training set) [13].

To identify a salient subgraph of the interactome to predict outcomes in large data sets and in parameter space smaller than the number of analytes, we propose the Graph Ensemble Neural Network (GENN), a novel type of GNN that learns a graph of linear analytes, associations predictive of both continuous (regressor-like) and discrete (classifier-like) outcomes. GENN produces an ensemble model of associations using a multi-layer pruning and pooling strategy, allowing it to make decisions about the saliency of associations and larger association groups that are connected within the graph. This new method reduces the number of parameters to train by computing *metafeatures* describing the interactome and then tuning only the weights associated with each metafeature, which are combined to obtain model weights for each edge in the graph (where nodes represent analytes). GENN further introduces two novel types of layers in its architecture: (**A**) *Composite Model Pooling* (CMP), a pooling procedure that rearranges the terms of the candidate predictors to obtain an interpretable function, and **(B)** *Significance Propagation* (SigProp), a pruning procedure that removes associations which do not improve outcome prediction. Using this approach, the number of parameters is equal to the number of metafeatures leading to a scalable yet robust method that can identify the optimal salient subgraph of the interactome. Figure 1 describes our overall method.

**Figure 1.**
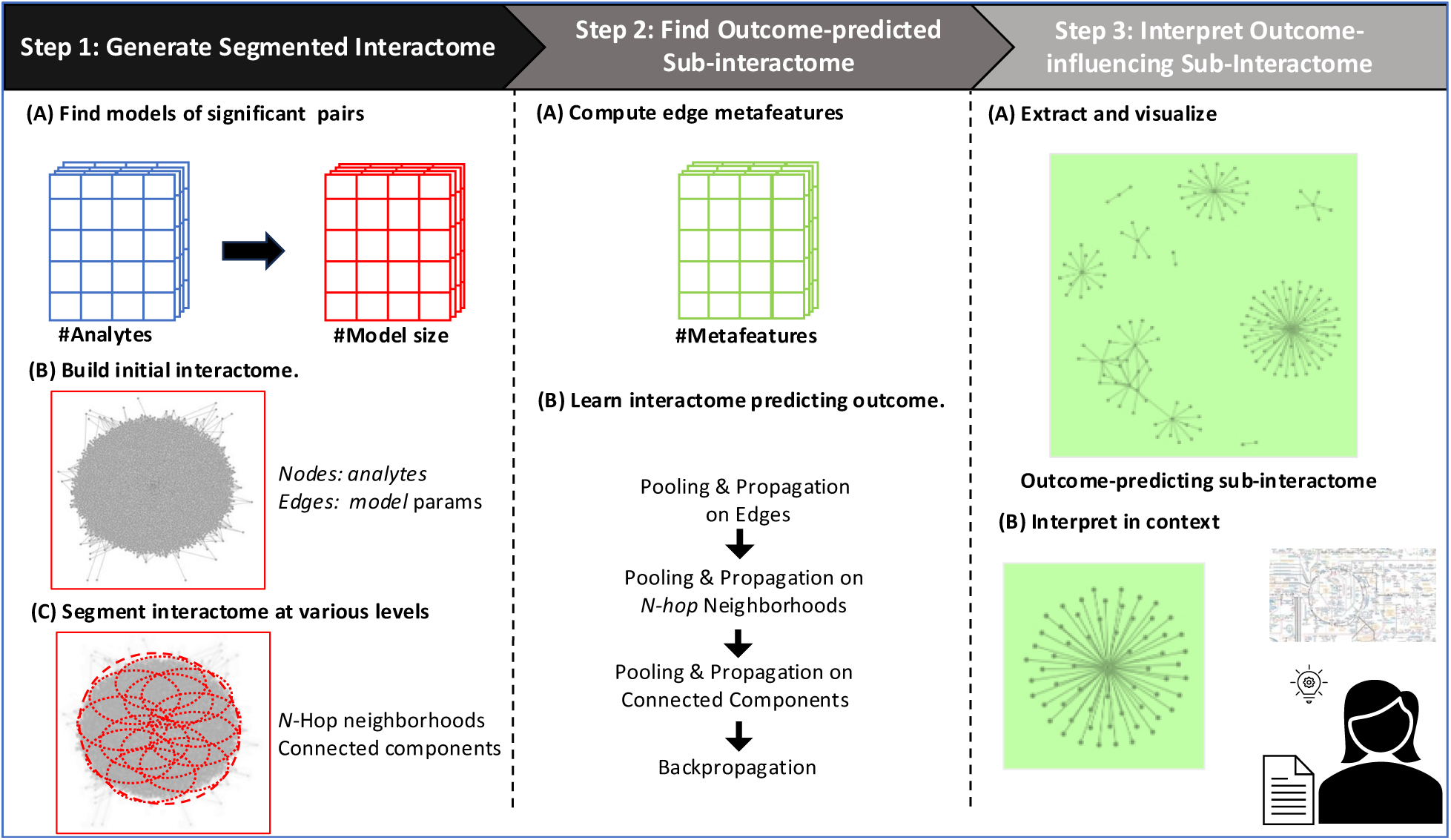
An outline of the procedure used by GENN to extract salient analyte subgraphs predictive of outcome.

We applied GENN to both theoretical models and to real data with matched transcriptomic and metabolomic measurements, to evaluate recovery of optimal analytes graphs. First, using theoretical data, we illustrate GENN’s ability to uncover ground truth analytes models, and compare the results to baseline models LASSO, Group LASSO, RF, and to multiple variants of GENN (which we refer to as *GNN + CMP* and *GNN + SigProp*) as an ablation study. Second, data from the National Cancer Institute Human Tumor Cell Lines Screen (NCI-60), TOPMed, and TCGA were evaluated using GENN. Using the NCI-60 data set, we illustrate GENN’s ability to uncover literature-supported analytes, pathways, metabolite classes, and gene sets predictive of outcome in real collected data. Then, using the TOPMed and TCGA datasets, we illustrate GENN’s ability to uncover potentially novel multi-omic associations driving total serum IgE in pediatric asthma and patient survival outcome in glioma, respectively.

Lastly, to show generality, GENN was applied to a subset of glioblastomas (GBM) (IDH wild type grade 4 astrocytomas) on available gene expression and DNA methylation data from TCGA. GENN discovered novel negative associations of various activation and killing mechanisms of innate immunity with worsening prognosis in GBM,

GENN offers several advantages over existing state of the methods. While current AI methods increasingly require large amounts of data and emphasize zero shot learning, we have created a method that extracts an interpretable interactome (and not in embedded space), scales well, is robust and can work with smaller amounts of data. Further, our method finds the sub-interactome that best influences outcome. Finally, GENN is generalizable and can work with analytes obtained from any of the omic regimens, include a multitude of metafeatures to ensure more robustness, and incorporate other measures of significance.

## Results

### 1. GENN is Robust to Noise Introduced in a Theoretical Model

We generated data sets with both continuous and discrete outcomes, two different analyte types with predefined outcome-dependent relationships, and noise applied to the dependent analyte type to evaluate GENN’s ability to exclude irrelevant analytes. We compared GENN results against other frequently applied methods and evaluated their ability to generate models including the correct analytes using a metric of our design termed the Analyte *F_1_* score (*AF_1_*) (**Methods**). As shown in **Table 1**, GENN was the best method to recover models that included the correct analytes when compared to other methods across varying noise levels., where the noise level corresponds to the variance of added Gaussian noise as a percentage of analyte variance.

**Table 1.** Results on the ability to exclude irrelevant analytes applied to theoretical data with noise.

|  | Noise Level | GENN | GNN + SigProp | RF | LASSO | Group LASSO | BN |
| --- | --- | --- | --- | --- | --- | --- | --- |
| Continuous | 1% | **0.97**<br>**(0.09)** | 0.69<br>(0.09) | 0.40<br>(0.10) | 0.36<br>(0.01) | 0.00<br>(0.00) | 0.43<br>(0.15) |
|  | 10% | **0.93**<br>**(0.13)** | 0.77<br>(0.15) | 0.40<br>(0.10) | 0.36<br>(0.01) | 0.00<br>(0.00) | 0.43<br>(0.13) |
|  | 50% | **0.86**<br>**(0.15)** | 0.67<br>(0.14) | 0.37<br>(0.12) | 0.36<br>(0.01) | 0.00<br>(0.00) | 0.40<br>(0.18) |
| Discrete | 1% | **0.89**<br>**(0.16)** | 0.57<br>(0.29) | 0.44<br>(0.14) | 0.36<br>(0.01) | 0.00<br>(0.00) | 0.66<br>(0.12) |
|  | 10% | **0.82**<br>**(0.18)** | 0.50<br>(0.28) | 0.42<br>(0.11) | 0.36<br>(0.01) | 0.00<br>(0.00) | 0.67<br>(0.13) |
|  | 50% | **0.64**<br>**(0.19)** | 0.36<br>(0.20) | 0.39<br>(0.13) | 0.36<br>(0.01) | 0.00<br>(0.00) | 0.66<br>(0.12) |

Additionally, GENN outperformed baselines in predictive performance for theoretical models with both continuous and discrete outcomes (**Supplementary Tables 1-2**). Notably, for the discrete outcome model with 50% noise, the F_1_ score was higher in GENN than in BN although the AF_1_ score was lower. Data simulation and hyperparameter selection are described in the **Supplementary Methods**.

### 2. GENN Improves Predictions and Ability to Recover Known Compound-Gene Relationships in NCI-60 Data

We evaluated GENN’s ability to extract salient interactomes from small sample sizes using a leave-one-out cross validation (LOOCV) approach in the NCI-60 metabolomic and transcriptomic data. The outcome was response to the chemotherapeutic drug Tegafur (class of 5-fluorouracils – 5-FU), and cancer type was included as a covariate. We evaluated the ability of GENN, baseline, and various ablation models to recover analytes and analyte groups known to be involved in 5-FU metabolism using a metric which we derived called *Reliability* (see **Methods**), which ranges from 0 to 1 where a higher score means higher reliability. Further, we evaluated the ability of all methods to build models of Tegafur metabolic response that predict correctly across a range of true responses using our *Scaled Covariance* (*SCov)* metrics defined in **Methods**, which ranges from - 1 to 1 where a higher magnitude means higher covariance and the sign corresponds to direction of covariance. For both *Reliability* and *SCov*, GENN outperformed other methods (**Table 2**). Pathways, gene sets, and compound sets used to compute the Reliability metric are defined in **Supplementary Tables 3-5**.

**Table 2.** Reliability and predictive performance of GENN models compared to other models to predict Tegafur response from NCI-60 transcriptomic and metabolomic data.

| Metric | GENN | GNN +<br>CMP | GNN +<br>SigProp | RF | LASSO | Group<br>LASSO |
| --- | --- | --- | --- | --- | --- | --- |
| Reliability | 0.19 | -- | 0.01 | 0.03 | 0.01 | 0.01 |
| SCov | 0.26 | 0.12 | 0.01 | -0.03 | -0.05 | -0.08 |

Notably, Reliability was not measured on the GNN + *CMP* model because this model does not prune the graph. GENN models yielded 4,521 gene-compound pairs included in at least one-fold, of which 14 pairs were shared across at least 10 LOOCV folds with consistent directionality. This result indicates that the relationship between the gene and compound is consistently positive or negative, suggesting robustness despite the small sample size. (**Supplementary Table 6**).

### 3. GENN Finds Novel Associations Between Sphingolipids, Serine and Threonine Metabolism, Immune Response, and Mitochondrial Organization Driving Total Serum IgE in Pediatric Asthma

We applied GENN to metabolomic and transcriptomic data from the TOPMed Childhood Asthma Management Program (CRA_CAMP) and Genetics of Asthma in Costa Rica Study (GACRS) pediatric cohorts to uncover multi-omics interplay underlying increased total serum IgE in pediatric asthma (outcome). Patient sex was used as a covariate. The graphs learned by GENN included 94 gene-compound pairs in CRA_CAMP (**Supplementary Table 7**) and 485 gene-compound pairs in GACRS (**Supplementary Table 8**), with 20 genes and 1 compound shared between GACRS and CAMP (**Supplementary Table 9**).

The composite graphs learned by GENN for the CRA_CAMP and GACRS TOPMed cohorts are shown in **Figure 2**.

**Figure 2.**
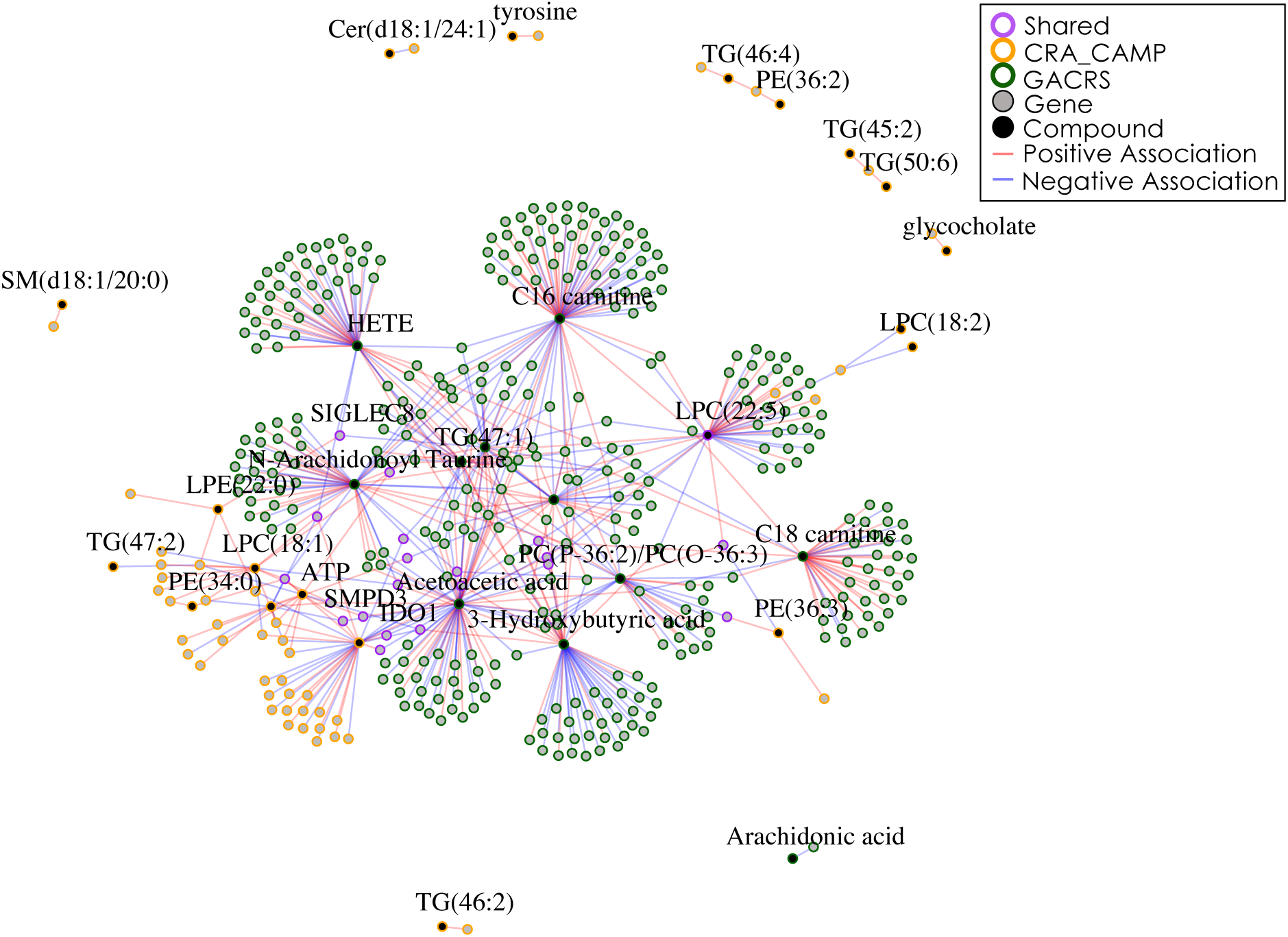
The combined graph resulting from GENN applied to two independent TOPMed cohorts. We observe large differences in the salient subgraphs predictive of total serum IgE in GACRS (outlined in green) and CRA_CAMP (outlined in yellow). However, several analytes overlap between the cohort-specific graphs (outlined in purple). Hub analytes from both graphs are labeled, with genes in gray and compounds in black.

Some of the analytes common to both TOPMed graphs were associated with literature-supported analyte groups and pathways from RaMP 2.0 [14] and GeneCards [15] (**Supplementary Table 9**), including amino acid metabolism (*IDO1*) [16, 17], apoptosis and immune response (*GZMB*, *CCR3*, *CEBPE*, *FBXW4*, *SIGLEC8*, *RAB44*, and *GPR114*) [16, 18–20], sphingolipid metabolism (*SMPD3*) [21–23], phosphatidylcholines (PCs) (LPC(22:5)) [24], and serine and threonine metabolism (*PRKY*). This graph suggests a possible interplay between these pathways in the context of total serum IgE in asthma. Despite these common analytes, investigation of the TOPMed cohort-specific graphs reveal a differential interactome between these pathways and other pathways (**Supplementary Table 10**). Analytes involved in inflammatory processes were associated with amino acids in CAMP, eicosanoids and fatty acids in GACRS [25, 26]. These results suggest that inflammation may be triggered by different molecule types in the CRA_CAMP and GACRS populations. Further, PCs were associated with the TCA cycle in GACRS, but with interleukins in CRA_CAMP, suggesting that PC may contribute to total serum IgE by multiple processes that are population-dependent.

Pathways (Holm-adjusted *p*-value < 0.05), metabolite classes (FDR-adjusted *p*-value < 0.05), and gene sets (*q*-values < 0.05) enriched for genes and compounds in the CRA_CAMP and GACRS predictive graphs include inflammatory analyte functional groups such as cytokine and chemokine pathways (**Supplementary Table 11**). Additionally, PCs were enriched in both cohorts, supporting their role as a key class of metabolites involved in inflammation. Conversely, eicosanoids, fatty acyls, and bile salts were only significant in GACRS, and imidazopyrimidines were only significant in CRA_CAMP, suggesting differing roles of these metabolite classes in the different populations. In summary, both the cohort-specific GENN models and the intersection of GENN models across cohorts highlights unique well-supported analytes in the context of total serum IgE in pediatric asthma (**Supplementary Table 12-13**).

### 4. GENN Finds Negative Associations Between JAK Kinase Activation and Killing Mechanisms of Innate Immunity with Worsening Prognosis in Glioblastomas

We applied GENN to a subset of glioblastomas (IDH wild type grade 4 astrocytomas) with available gene expression and DNA methylation data from TCGA [27] **(Supplementary Table 14)**. Patients were subset into two prognosis groups with differing survival outcomes (log-rank *p* < 0.01) using subspace clustering [28]. These groups were used as outcome in GENN. The resulting GENN model included a total of 38 expression-methylation associations **(Supplementary Table 15)**. This workflow is described in **Figure 3**.

**Figure 3.**
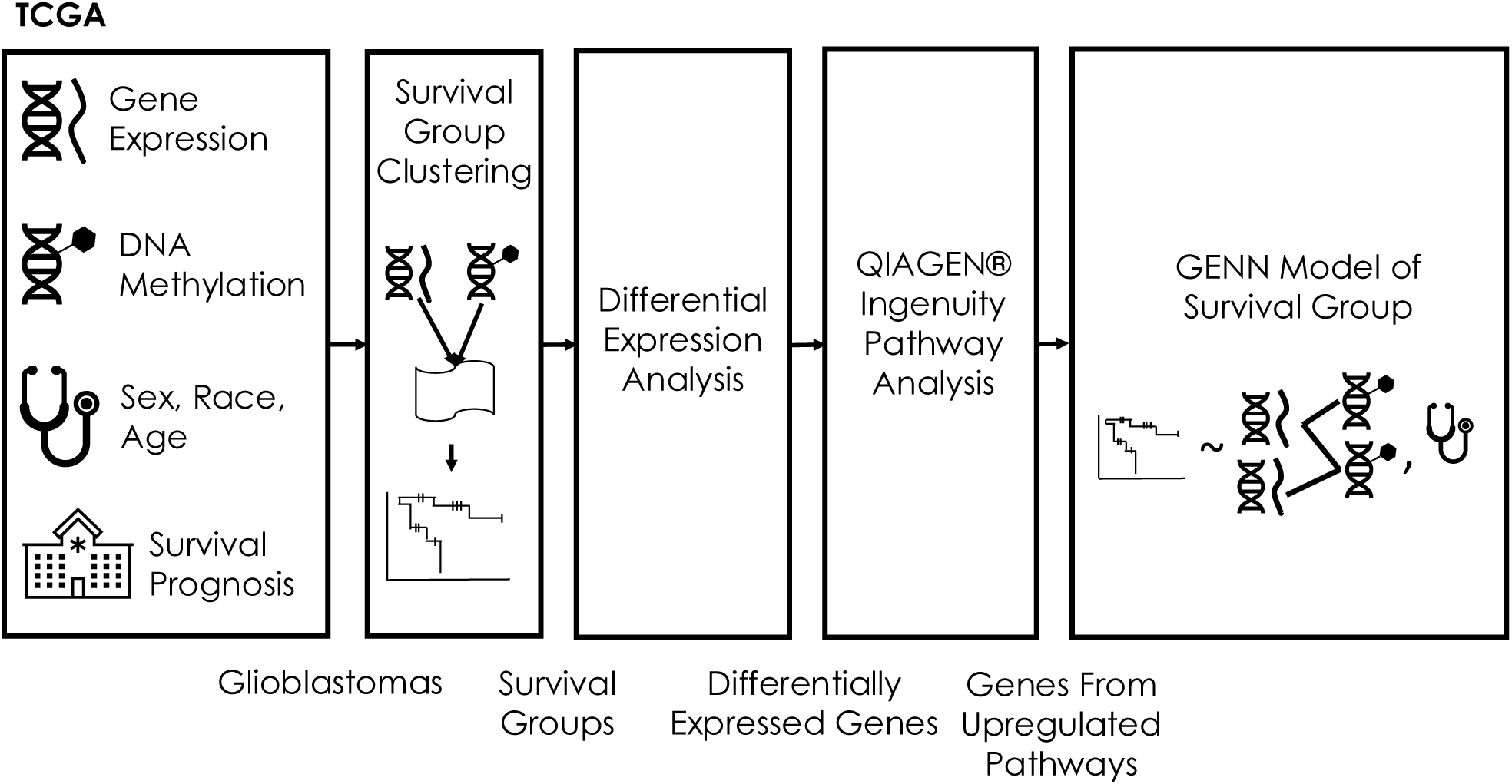
The analytical workflow for the TCGA data. Multi-omics data were input into a subspace clustering framework to obtain survival groups. Differential expression analysis was then performed on the genes and IPA run on the differentially expressed genes. All genes from upregulated pathways and all methylation probes from promoter regions were input into GENN, with sex, race, and age included as the covariates and the survival groups as the outcome.

The resulting graph is visualized in **Figure 4**.

**Figure 4.**
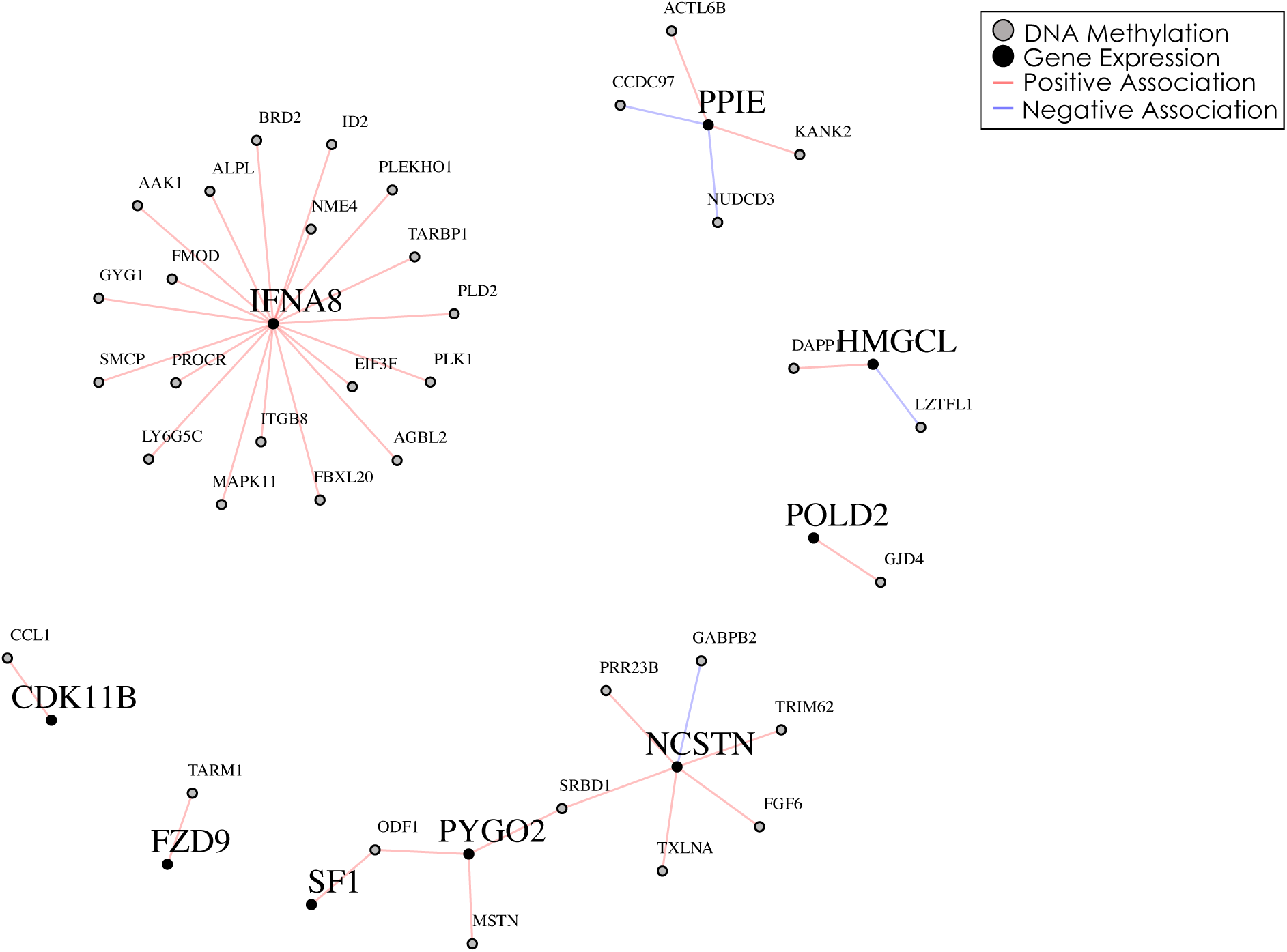
The graph resulting from GENN applied to TCGA glioblastoma data. We observe a hub of associations between IFNA8 and methylation probes that do not associate with other genes in the graph. Here, a positive association indicates that increased methylation of the source node corresponds to increased expression of the target node with respect to cluster. The graphs with smaller font indicate methylation levels.

Notably, a hub of positive associations (i.e., cluster-dependent associations between increased expression and increased methylation) was centered around expression of *IFNA8*, which includes deletions that have previously been associated with poor prognosis in gliomas with *CDKN2A/B* and *EGFR* alterations [29] and with single nucleotide polymorphisms (SNPs), such as *IFNA8* rs12553612 AC genotype, which was related to improved prognosis in glioma patients [30]. Further exploration of *IFNA8* and its association with methylation probes revealed that the associated probes were on different chromosomes from *IFNA8* **(Supplementary Table 16)**, suggesting that the methylation-expression interactome likely consisted not of trans-regulatory associations but rather associations between hypermethylated genes and *IFNA8*.

*IFNA8* encodes interferon *α*, which activates JAK kinases, known to promote treatment resistance in glioblastomas (Reactome: R-HSA-877336) [31, 32]. Among hypermethylated genes associated with *IFNA8* expression we found *GYG1*, which encodes glycogenin, discharged by neutrophils during exocytosis (Reactome: R-HSA-6801466) [33], and *PLD2*, which encodes phospholipase D and plays a positive role in phagocytosis (Reactome: R-HSA-2029485) [34]. These associations suggest, respectively, that both neutrophil degranulation (Reactome: R-HSA-6801466) [33] and phagocytotic activity (Reactome: R-HSA-2029485) [34] decrease as JAK/STAT signaling increases, therefore the tumor becomes more resistant to treatment.

Relatedly, *IFNA8* also signals *PIK3CD* encoding in the phosphoinositide 3-kinase (PI3K) complex in PI3K-Akt signaling and was associated with hypermethylation of *ITGB8*, which encodes an integrin subunit and signals PIK3CA encoding in the PI3K complex (WikiPathways: WP4172) [35], suggesting that the presence of PIK3DC and absence of PIK3CA in in the PIK complex may increase resistance to treatment.

Other associations of note include hypermethylation of *MAPK11*, which was positively associated with *IFNA8* expression in the context of worse prognosis. *MAPK11* encodes p38 mitogen-activated protein kinase (p38 MAPK), which can function as either a tumor suppressor or an oncogene depending on the tumor microenvironment [36]. Hypermethylation of *PROCR*, which encodes the receptor for protein C, resulting in downregulation of the protein C anticoagulant pathway, can result in venous thromboembolism [37, 38]. Patients with glioblastoma are at high risk for venous thromboembolism, and while this complication is not independently associated with survival [39], our results indicate that the combination of venous thromboembolism and JAK activation may result in worsening prognosis of glioblastoma patients. Finally, hypermethylation of *BRD2* forms a complex with *RUNX3* during the restriction point (R-point) to regulate the cellular transition [40].

This interactome provides insight into the mechanisms driving worse patient prognosis in glioblastomas, suggesting interplay between JAK kinase activation and the killing mechanisms of innate immunity as well as other pathways.

### 5. Running Time Analysis

**Supplementary Table 17** lists empirical running times for GENN, GNN + SigProp, and GNN + CMP on the theoretical models, NCI-60 data, and TOPMed data. Setup for theoretical models was run on a laptop running macOS 13.5 with a 6-Core Intel Core i7 processor, and all other analyses were run on a high-performance computing cluster running CentOS Linux 7 with Intel Gold 6140 CPU’s. Setup (i.e., construction of the graph and computation of metafeatures) was the same for all models.

## Discussion

GENN obtains salient, interpretable interactome subgraphs by using associations to build predictive models of outcome. In real data from multiple biomedical contexts, GENN models both **(i**) included reliable analytes and analyte groups supported by the literature, and (**ii**) predicted outcome more reliably than comparable methods. Further, GENN is robust to varying degrees of Gaussian noise added to the analyte measurements in theoretical models. In contrast, other methods appear to converge on locally optimal solutions rather than isolating the correct set of analytes. This is particularly true of LASSO and Group LASSO, whose reduced performance can likely be explained by the fact that, in the theoretical models, the outcome is not a linear function of the analyte measurements.

GENN models trained on the TOPMed asthma cohorts suggested common roles of serine and threonine metabolism and the downstream TCA cycle as well as PC metabolism pathways, echoing the results of Eicher et al [41]. Further, immune signaling has been previously linked with tubulin [42], choline [43], and hyaluronic acid [44], as reflected in the enrichment of tubulin modification and hyaluronan metabolism in CRA_CAMP. Additionally, previous work has discussed the roles of eicosanoids [45] and PCs [46] on neutrophil degranulation and the role of bile acids in altering the crystallization of cholesterol in the presence of sphingomyelin [47, 48], as reflected in the crosstalk between eicosanoids, PC, and neutrophil degranulation and between bile salts and sphingolipids in GACRS. Finally, the associations between amino acids and inflammatory processes reflect recent work by Li et al. showing that ferroptosis (which is regulated by amino acid metabolism) triggers inflammation in asthma [49]. We note that the differences in analyte crosstalk and pathway enrichment between the TOPMed cohorts are expected because the CRA_CAMP and GACRS cohorts differ in several respects, namely: (**i**) the sample size of CRA_CAMP is larger than that of GACRS, (ii) CRA_CAMP patients are older (and relatedly, exhibit higher weight and height measurements) than GACRS patients on average, (**iii**) environmental and dietary metabolite exposures are expected to differ due to geographical differences, and (iv) the CRA_CAMP and GACRS patients have different ethnicities.

Furthermore, GENN models trained on data from TCGA suggested that concordant upregulation of the JAK-STAT pathway and downregulation of innate immune killing responses, in combination with venous thromboembolism risk and R-point modifications, are associated with worse prognosis in glioblastoma. Moreover, while previous work investigating the PI3K catalytic subunit function on glioblastoma did not find associations between these subunits and glioblastoma cell survival independently [50], we are not aware of any research that has investigated them in tandem. Furthermore, activation of the PI3K complex has been postulated to increase phospholipase D [34], supporting the negative association between phagocytotic activity and JAK activation with worsening prognosis in glioblastoma.

We note that GENN differs from a traditional GNN in multiple ways. While a traditional GNN applies neural network layers of the graph, the edges of the GENN graph represent candidate predictors that are combined for prediction using an ensemble method applied to each candidate-pair predictor and its metafeatures, as introduced by Cruz et al. [51]. We devised metafeatures appropriate for a graph learning context and used them to calculate model weights. Additionally, while a traditional GNN may include convolution and pooling layers, we introduced novel types of layers which are integral to GENN.

We note that related work by Ding et al., known as cooperative learning, also builds a multi-omics ensemble model wherein model consensus is optimized [52]; however, cooperative learning does not model an interactome between multi-omics data. Moreover, unlike BN, discovery of the interactome using GENN is a tractable problem for large multi-omic data sets. Because GENN outputs a graph of the underlying interactome driving outcome, the model can be easily interpreted in the context of known literature and known functional relationships between the analytes.

There are several limitations and directions for future improvement of GENN. First, the metafeatures described herein represent a small subset of possible metafeatures that could be used in training GENN. Moreover, metafeatures are specific to each analyte pair, and the current work therefore does not leverage performance metrics or other types of metafeatures at additional levels of the graph (e.g., 1-hop neighborhood-specific metafeatures), which could be beneficial for improving model performance. Model performance may be further improved using different types and levels of graph segmentation (see **Supplementary References #6**). Additionally, running time and memory constraints may be prohibitive for very large sample sizes and analyte counts; this could be improved by developing pruning heuristics preventing segmented graph neighborhoods from overlapping and pruning in only a subset of training iterations. We also note that the evaluations in this work focused on two -omics data types in each evaluation scenario (either transcriptomics and metabolomics or transcriptomics and methylomics). However, we hypothesize that GENN can readily be expanded to evaluating pairwise associations between three or more - omics data types. To do so, GENN could be expanded to perform independent runs of IntLIM 2.0 on each pair of data types, which would then be consolidated in tandem using the same procedure as is used with two data types.

We believe that the real value of GENN and its derivatives will be apparent when data sizes are much larger (viz. biobanks), when finding salient associations from billions is essential, although appropriate scaling of the methodology will be required for this type of application. Additionally, GENN-like methods can help with the design of experiments; smaller batches can be used to stratify the underlying subject before launching collections-at-scale.

## Methods

### Overview of GENN

To find subgraphs of the interactome that predict sample outcomes, GENN first generates an analyte association graph, where nodes are analytes and edges are associations, using linear outcome-dependent association models for each pair of analytes. GENN then finds an optimal subgraph predictive of outcome by (1) pruning the graph using significance propagation (*SigProp*) and composite model pooling (*CMP*) perceptrons, and (2) optimizing weights associated with interactome metafeatures (see Methods). Essentially, GENN learns the subgraph of differential correlation models and hence analyte pairs with least errors and most effect size on the outcome. Once the predictive model is learned, users can then interpret it in the context of known literature and functional annotations.

### Model Formulation

GENN learns a predictive graph *G* where nodes *N(G)* represent features (analytes) in feature sets *S_1_,…,S_𝔐_*

corresponding to -omics data types 1,…, 𝔐 respectively, and edges *E(G)* represent linear outcome-dependent associations between the nodes corrected for an optional set of covariate features *S_𝔐+1_*. The data input to GENN includes the following matrices and vectors:

- For 1 ≤ *i* ≤ 𝔐, measurement matrices *X*^S_i_^ with dimensionality *n* × |*S_i_*|, where *n* is the number of samples in a data set and 𝔐 is the number of omics data types.
- An optional covariate feature matrix *X*^*S*_*𝔐*+1_^ with dimensionality *n* × |*S*_𝔐+1_|.
- An outcome vector *Y* with dimensionality *n*.

In this work, we applied GENN to data sets with 𝔐 = 2. GENN obtains a linear model for each pair of analytes using the *IntLIM 2.0* R package [53] and described in **Equation 1**, where *i* and *j* are analyte indices, *X_c_* is the input vector for covariate *c*, and |*S*_3_| is the number of total covariates.

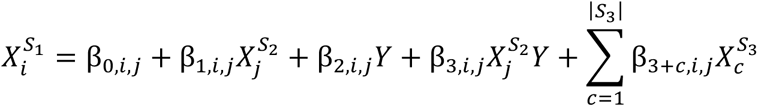

**Equation 1**. Definition of edges E(G) between analytes S_1_ and S_2_.

Further, we define a tensor of *Γ metafeatures M* with dimensionality *n* × |*E*(*G*)| × Γ. Each metafeature measures the strength of an interaction or its predictive ability in a different manner, to simplify the parameter space. The rationale for defining metafeatures in this way is that we expect the most salient interactome to include the associations that either predict outcome well independently or represent statistically significant outcome-dependent associations. In this work, Γ = 5, and the set of metafeatures is described in the **Supplementary Methods**. Given *G*, *M*, and *ß*, GENN obtains a composite predictor of outcome as defined in **Equation 2**, where *k* is the sample index.

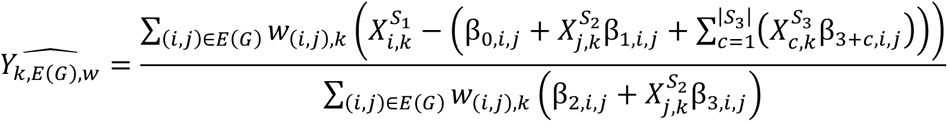

**Equation 1. GENN composite predictor.**

Weights w_(i,j),k_ are defined in **Equation 3**, where *ϕ* is the learned weight of a metafeature (*ϕ_γ_* for metafeature *γ*). Learning only Γ parameters instead of |S_1_ × S_2_| drastically reduces the model parameter space.

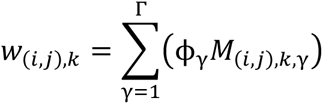

**Equation 2. Formula for computing edge weights in GENN.**

### Optimization Function and Training Procedure

The optimization function is defined in **Equation 4**.

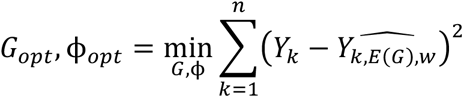

**Equation 3. GENN optimization function.**

We estimate *G_opt_* using a combination of pooling at the 1-hop neighborhood, connected component, and whole graph levels (Composite Model Pooling or *CMP*) and a pruning procedure that propagates only edges that reduce the distribution of model error (Significance Propagation or *SigProp*) (**Figure 5**).

**Figure 5.**
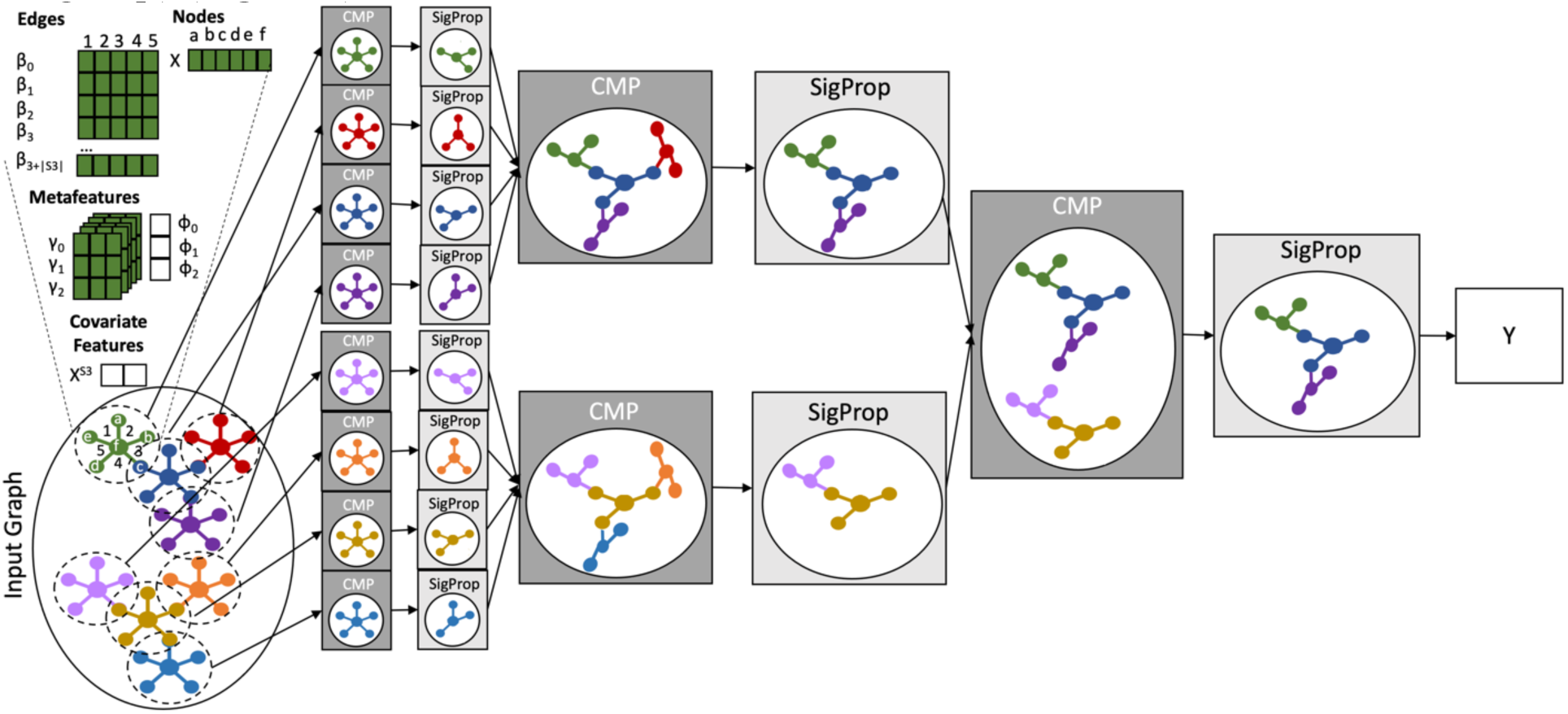
GENN architecture. The inputs to GENN include an input graph, sample-specific metafeatures, initialized metafeature weights, covariate features, node annotations, and edge annotations. The architecture consists of three layers of Composite Model Pooling (CMP) and Significance Propgation (SigProp), corresponding to 1-hop neighborhoods, connected components, and the whole graph. In each stage of training, subgraph-specific models are pooled using CMP and pruned using SigProp.

We optimize *ϕ* using backpropagation with the ADAM optimizer [54] for the theoretical models, NCI-60 data, and TOPMed data and Stochastic Gradient Descent for the TCGA data (**Supplementary Algorithm 1**). The gradient for the optimization function with respect to *ϕ* is given in **Equation 5**, where *ε* is the model error, and *G_ι_* and *ϕ_ι_* represent the graph and metafeature weights for training iteration *ι*.

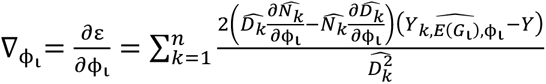

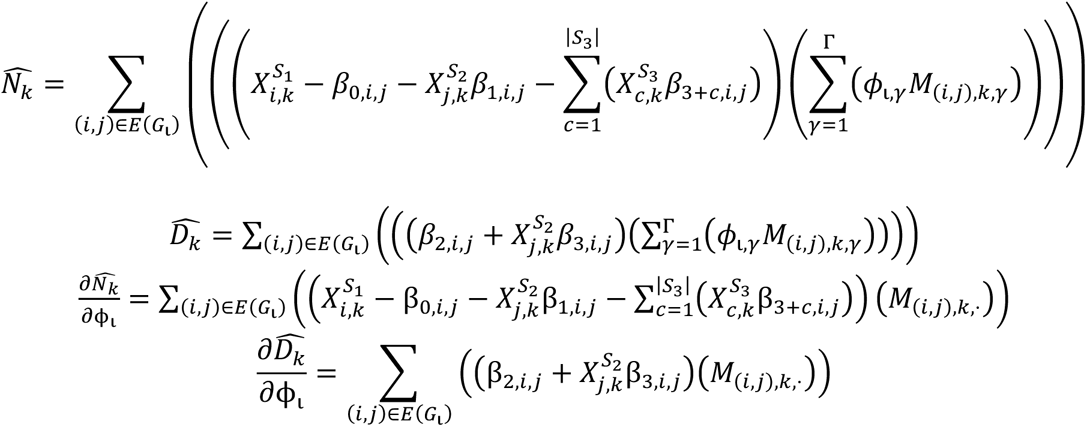

**Equation 4. Gradient for training GENN.**

### Composite Model Pooling (CMP)

In the first layer of the graph (*ℓ* = 1) edges (*H*_0_* = *E(G)*) are combined into 1-hop neighborhoods (*H_1_*) (**Supplementary Algorithm 2**). In the second layer (*ℓ* = 2), pruned 1-hop neighborhoods (*H*_1_*) are combined into connected components (*H_2_*) found using breadth-first search. In the third layer (*ℓ* = 3), pruned connected components (*H*_2_*) are combined at the whole graph level (*H_3_*). Local predictors of outcome 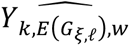 are constructed for each edge, neighborhood, or connected component index *ξ* in layer *ℓ*.

### Significance Propagation (SigProp)

For a given component index *ξ* in layer *ℓ*, the significance of a subgraph *G_ξ,ℓ_*[*ζ*] in the context of *G_ξ,ℓ_* is computed by comparing t(*G_ξ,ℓ_*, *ϕ, Y*) to t(*G_ξ,ℓ_* - *G_ξ,ℓ_*[*ζ*], *ϕ, Y*), where *t* is a variation of Welch’s *t*-statistic given in **Equation 6**.

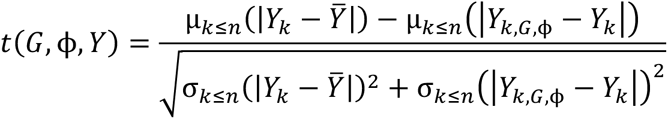

**Equation 5. The significance of a subgraph in GENN.**

Subgraphs are greedily pruned using this metric (**Supplementary Algorithm 3**).

### Analyte F_1_ Score

To evaluate a model’s inclusion of analytes from ground truth predictive associations (*V_sim_*), we introduce *AF*_1_ score as defined in **Equation 7**, which is derived from the *F*_1_ classification metric, where *V_opt_* is the set of analytes included in *G_opt_*.

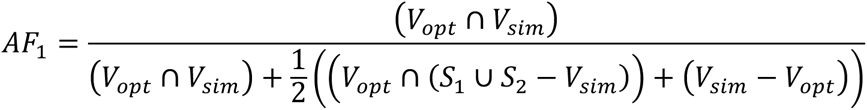

**Equation 7. Formula for Analyte F1 score.**

### Scaled Covariance Metric

In unevenly distributed data, some methods yield predictions that are biased and limited to a certain range [55]. To measure the ability of each method to isolate salient analyte groups that predict continuous outcomes across the expected outcome range when data are unevenly distributed, we introduced a predictive correlation metric adjusted for range in unevenly distributed data known as *Scaled Covariance* (*SCov*) (**Equation 8**).

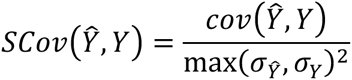

**Equation 8. Formula for scaled covariance.**

*SCov* accounts for predictor range in a manner other regression metrics do not, as illustrated in the **Supplementary Methods**.

### Reliability

For experimental data, we assessed the reliability of the graph using literature support for individual analytes (genes and metabolites) and analyte groups (biological pathways, gene sets, and metabolite classes) as described in **Equation 9** where *V*_O"9_is the set of analytes and analyte groups supported by the literature, and *V*_789_ is the set of analytes and analyte groups included in *G*_789_. This metric is asymmetric because a lack of literature support does not imply that a result is invalid; it simply may have not been discovered yet.

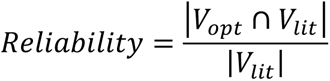

**Equation 9. Formula for model reliability.**

### Study Design

Due to the inherent challenges of establishing ground truth analytes and analyte sets (e.g., gene sets and pathways) underlying outcome in real data, we evaluated GENN’s ability to uncover ground truth using noisy theoretical models of continuous and discrete outcome (**Supplementary Methods**). We measured the ability to recover the true model using the Analyte *F*_1_(*AF*_1_) score. described further in the **Supplementary Methods**. We measured predictive performance on the theoretical data sets using *SCov* for continuous outcomes, and the F_1_ Score, True Positive Rate (TPR), and False Positive Rate (FPR) for discrete outcomes. Using these metrics, we compared GENN against the following baselines: (**i**) LASSO, (**ii**) RF, (**iii**) Group LASSO, and (**iv**) BNs, where the groups are the 1-hop neighborhoods of the interactome. In addition, as an ablation study, we compared GENN to (**iv**) GNN + CMP, a variation of GENN that excludes *SigProp*, and (**v**) GNN + *SigProp*, a variation of GENN that uses predictor averaging to pool associations instead of using *CMP*. In theoretical models, BN were also included as a baseline. As an ablation study, two variants of edge-based graph prediction GNN were evaluated on both theoretical models and real data sets: GNN + Composite Model Pooling (GNN + CMP) and GNN + *SigProp* Pruning (GNN + *SigProp*).

We further evaluated the performance of GENN on the NCI-60 data set (**Table 3**). Because cancer cell response to 5-FU drug families, such as Tegafur, varies across cancer types [56], cancer type was used as a covariate both in GENN and other models evaluated. We evaluated performance across LOOCV folds using *SCov*. To measure each method’s ability to recover reliable analytes from the literature, we manually compiled literature relevant to each outcome prior to learning any models and devised a metric known as *Reliability*, which is based on manually curated knowledge from corpora of literature specific to 5-FU uptake mechanisms detailed in the **Supplementary References #8**, which was compiled as described in the **Supplementary Methods**.

**Table 3.** Characteristics of all biomedical data sets to which GENN was applied.

| Database | Human Tumor Cell Lines Screen | TOPMed<sup>1</sup> |  | TCGA<sup>2</sup> |
| --- | --- | --- | --- | --- |
| Data Set | NCI-60 | CRA_CAMP<sup>3</sup> | GACRS<sup>4</sup> | Glioblastoma Multiforme |
| Description | Cancer cell assay for drug response | Pediatric asthma cohort (North America) | Pediatric asthma cohort (Costa Rica) | Glioblastoma |
| Study Type | In vitro | Clinical trial | Observational study | Mixed |
| Number of Genes | 17,987 | 22,772 | 22,772 | 402 |
| Number of Named Compounds | 109 | 529 | 529 | NA |
| Number of Methylation Probes | NA | NA | NA | 122,849 |
| Number of Samples | 51 cell lines | 564 patients | 309 patients | 121 samples |
| Model Covariates | Cancer type | Patient sex | Patient sex | Patient sex, Patient age, Patient race |
| Outcome | Tegafur response | Total Serum IgE Level | Total Serum IgE Level | Prognosis-associated cluster |
| Program | Human Tumor Cell Lines Screen | National Heart, Lung, and Blood Institute Trans-Omics for Precision Medicine (TOPMed) | National Heart, Lung, and Blood Institute Trans-Omics for Precision Medicine (TOPMed) | TCGA |
| References | [57, 58] | [59-61] | [59-61] | [27] |

1 - TOPMed: Trans-Omics for Precision Medicine program. 2- The Cancer Genome Atlas. 3- CRA\_CAMP: The Genetic Epidemiology of Asthma in Costa Rica and the Childhood Asthma Management Program. 4- GACRS: The Genetics of Asthma in Costa Rica Study

To evaluate GENN’s ability to uncover novel mechanisms underlying disease or phenotype using different types of multi-omic data, we applied GENN to two studies from TOPMed, where the outcome is total serum IgE level, and to a biomedical data set from TCGA, where the outcome is prognosis-associated cluster in glioblastoma (**Table 3**). In the TOPMed cohorts, sex of subjects was used as a covariate. In the TCGA data, sex, age, and race of subjects were used as covariates. GENN was trained using all DNA methylation probes from promoter regions of protein coding genes and all expression levels from pathways upregulated in the cluster with worse survival outcome using QIAGEN IPA (QIAGEN Inc., https://digitalinsights.qiagen.com/IPA).

### Data Pretreatment

Data pretreatment for TOPMed asthma cohorts is described in Eicher et al [41]. NCI-60 data transcriptome and metabolome measurements were obtained from Su et al. [58], and the IC_50_ drug score matrix was downloaded from CellMiner [57]. Drug score was drawn from experiment 9403SS94, which had a wider range of cell-line-wise means than the other 15 experiments including the 5-FU drug family (**Supplementary Figure 1**). Glioblastoma data were downloaded from TCGA, and race, sex, and age at diagnosis drawn from TCGA sample files. Expression levels were transformed using the *VOOM* R package [62]. Methylation probes with NA values for all samples were removed, and the remaining NA probes imputed using a custom parallelized version of the *MethyLImp* R package [63].

### Baselines and Ablation Study

Baselines were evaluated using the R *randomForest* [64], *glmnet* [65], *gglasso* [66], and *bnlearn* [67] packages for RF, LASSO, Group LASSO, and BN, respectively, and the groups in Group LASSO were defined as *H_1_*. The ablation study is described further in the **Supplementary Methods**, with GNN + CMP training described in **Supplementary Algorithm 4**.

## Supporting information

Supplementary References

## Data Availability

Data are available from the authors upon request.

## Code Availability

GENN can be downloaded and installed from https://github.com/machiraju-lab/GENN. Scripts to replicate results are located in https://github.com/machiraju-lab/GENN_Paper_Scripts.

## Acknowledgments

We thank the NIH Individual Graduate Partnership Program for training and mentoring. Further, we thank the Broad Institute for their correspondence regarding the metabolomics data. The Genetics of Asthma in Costa Rica Study and the Childhood Asthma Management Program were supported by grant HL132825 from the National Institutes of Health. Metabolomic analyses were supported by R01HL123915, R01HL155742, 1R01HL141826, and W81XWH-17-1-0533. Further NIH support comes from R01HL155742. This research was supported in part by the National Center for Advancing Translational Sciences (NCATS) Informatics Research Core (ZIC TR000410) and the National Cancer Institute [R01CA169368, R01CA11522358, R01CA1145128, R01CA108633, R01CA188228, 1RC2CA148190, and U10CA180850-01], A Brain Tumor Funders Collaborative Grant, and the Ohio State University Comprehensive Cancer Center Award.

## Author Contributions

EAM, RM, and TE conceptualized the study. TE developed the GENN package and completed the analyses of the theoretical models and real data. STW, RG, MM, JC, RSK, and JL-S contributed to the design and collection of the TOPMed data described herein. CC performed metabolomic profiling for the TOPMed data described herein. AB, JH, JM, WM, and AC contributed to the design of the TCGA workflow. AB annotated the TCGA data described herein. TE, AW, and JM preprocessed the TCGA data described herein. RSK, JL-S, EAM, AB, JH, JM, WM, TE, and JC contributed to data interpretation. TE, EAM, AB, JC, and RM drafted the manuscript and contributed to revision of manuscript and editing. All authors had final responsibility for the decision to submit for publication.

## Competing Interests

STW receives royalties from UpToDate Inc. and is on the Board of Histolix Inc. JL-S is a scientific advisor to TruDiagnostic Inc, Precion Inc, and Ahara Corp and is on the Metabolomics Society Board.

## Materials and Correspondence

Correspondence and material requests should be addressed to RM.

