## Supplementary Methods for "Finding Salient Multi-Omic Interactomes Relevant to Multiple Biomedical Outcomes using Graph Ensemble Neural Networks"

### Supplementary Algorithms

**Data:** $X$, $Y$, learning rate η, Γ, iterations $I$, convergence cutoff $d$, layer count $L$
**Result:** $G_{opt}, \phi_{opt}$
$\beta, G_0, H, M, \phi_0 \leftarrow Initialize(X, Y, \Gamma)$
// Train until convergence.
**While** $\iota < I$ & $\delta < d$ **do**
    $H_1^* \leftarrow E(G_0)$ // In layer 1, each edge is a subgraph.
    **For** $l \in Set(2, \ldots, L)$ **do**
        **For** $\xi \in Set(1, \ldots, |H_l|)$ **do**
            // Pool and prune mapped subgraphs from the previous layer.
            $H_l^*[\xi] \leftarrow SigProp(Pool(H_l[\xi], H_{l-1}^*), \phi_{\iota-1}, M, \beta, Y)$
        **end**
    **end**
    $G_\iota \leftarrow \cup (H_L^*)$ // Update graph.
    $\phi_\iota \leftarrow ADAM\left(\eta, \nabla_\phi(G_\iota, Y, X, \beta, M, \phi_{\iota-1})\right)$ // Update metafeature weights.
    $\delta \leftarrow |\phi_{\iota-1} - \phi_\iota|, \iota \leftarrow \iota + 1$ // Update delta and iteration.
**end**
$\phi_{opt} \leftarrow \phi_\iota, G_{opt} \leftarrow G_\iota$ // Obtain final results.

Supplementary Algorithm 1. GENN training procedure. Initialization includes: (**i**) learning β using IntLIM 2.0, (**ii**) generating $G$ and partitioning at multiple levels to obtain $H$, and (**iii**) initializing ϕ and computing $M$.

**Data:** Graph segmented into edges $H_1$
**Result:** Graph segmented into neighborhoods $H_2$
$Q \leftarrow E(G), H\_2 \leftarrow \emptyset, \xi \leftarrow 0$ // Add all edges to a queue.
// Create new neighborhoods until queue is empty.
**While** $|Q| > 0$ **do**
    $(i,j) \leftarrow Q[1]$ // Select an edge from the queue.
    // The neighborhood is the edge and all adjacent edges
    $H_2[\xi] \leftarrow H_2[\xi] \cup Set((i,b) \in E(G)) \cup Set((a,j) \in E(G))$ // Build neighborhood.
    $Q \leftarrow Q - H_2[\xi]$ // Dequeue the neighborhood.
    $\xi \leftarrow \xi + 1$ // Start a new neighborhood.
**end**

Supplementary Algorithm 2. Neighborhood segmentation for GENN.

**Data:** unpruned subgraph $G_\xi$, $\phi$, $Y$
**Result:** pruned subgraph $H_l^*[\xi]$
$G_1 \leftarrow G_\xi, G_2 \leftarrow \emptyset$ // Initialize forward and backward pruning.
// Prune over all subgraphs from the previous layer mapping to the current layer.
**For** $G_\zeta \in G_\xi$ **do**
    // Backward prune if the criteria are met.
    **If** $t\left(G_1 - G_\zeta, \phi, Y\right) < t(G_1, \phi, Y)$ **do**
        $G_1 \leftarrow G_1 - G_\zeta$
    **end**
    // Forward prune if the criteria are met.
    **If** $t\left(G_2 \cup G_\zeta, \phi, Y\right) < t(G_2, \phi, Y)$ **do**
        $G_2 \leftarrow G_2 \cup G_\zeta$
    **end**
    // Choose between the forward pruned and backward pruned subgraph.
    $H_l^*[\xi] \leftarrow G_1$
    **If** $t(G_1, \phi, Y) < t(G_2, \phi, Y)$ **do**
        $H_l^*[\xi] \leftarrow G_2$
    **end**
**end**

Supplementary Algorithm 3. SigProp for subgraph $G_{\xi,\ell}$, in GENN.

**Data:** $X$, $Y$, learning rate η, Γ, iterations $I$, convergence cutoff $d$
**Result:** $\phi_{opt}$
$\beta, G, H, M, \phi_0 \leftarrow Initialize(X, Y, \Gamma)$
// Train until convergence.
**While** $\iota < I$ & $\delta < d$ **do**
    $\phi_\iota \leftarrow ADAM(\eta, \nabla_\phi(G, Y, X, \beta, M, \phi_{\iota-1}))$ // Update metafeature weights.
    $\delta \leftarrow |\phi_{\iota-1} - \phi_\iota|, \iota \leftarrow \iota + 1$ // Update delta and iteration.
**end**
$\phi_{opt} \leftarrow \phi_\iota$// Obtain final results.

Supplementary Algorithm 4. GNN + CMP training procedure

### Metafeatures

Each metafeature value for an edge is computed as a percentile $Q$ with respect to all other edges. Because weights $w$ are irrelevant for single-edge predictors, $w = 1$.

1. **Predictor consensus** quantifies the consensus of a predictor obtained from the single association $(i, j)$, i.e., $\widehat{Y_{k,(\iota,J),w}}$, with predictors obtained from all other edges in $E(G)$. It is defined in Supplementary Equation 1, where $Pr_{(a,b)\in E(G)}$ is the probability distribution over all edges in $E(G)$.

$$M_{i,j,k,1} = Q_{E(G)}\left(Pr_{(a,b)\in E(G)}\left(\widehat{Y_{k,(\iota,\jmath),w}} = \widehat{Y_{k,(a,b),w}}\right)\right)$$

Supplementary Equation 1. Formula for the prediction consensus metafeature.

2. **Outcome-dependent interaction *p*-value** is the significance of outcome-dependent interaction between entities $i$ and $j$, measured as a $p$-value from a linear regression $t$-test and defined in Supplementary Equation 2. $T$ is a $t$-distribution with $n-2$ degrees of freedom, $H_0$ is the null hypothesis, and $B_{3,i,j}$ is the true value of $\beta_{3,i,j}$ over a universal population [1].

$$M_{i,j,k,2} = Q_{E(G)}\left(1 - Pr\left(|T| \geq |\beta_{3,i,j}|, \operatorname{sgn} T = \operatorname{sgn} \beta_{3,i,j}\right) \middle| H_0 : B_{3,i,j} = 0\right)$$

Supplementary Equation 2. Formula for the outcome-dependent interaction $p$-value metafeature.

3. **Outcome-dependent interaction effect size** also measures the significance of outcome-dependent interaction between entities $i$ and $j$, but using a statistical effect size rather than a $p$-value. It is defined in Supplementary Equation 3.

$$M_{i,j,k,3} = Q_{E(G)}\left(|\beta_{3,i,j}|\right)$$

Supplementary Equation 3. Formula for the outcome-dependent interaction effect size metafeature.

4. **Interaction effect size** measures the significance of interaction that is not outcome-dependent, defined in Supplementary Equation 4.

$$M_{i,j,k,4} = Q_{E(G)}\left(|\beta_{1,i,j}|\right)$$

Supplementary Equation 4. Formula for the interaction effect size metafeature.

5. **Local error** measures the absolute error of the predictor on the $K$ nearest neighbors of sample $k$ $knn(k)$ in the training set, defined in Supplementary Equation 5. To find the $K$ nearest neighbors across multiple types of entity measurements, $k$ and other samples in the training set are projected onto a Grassmann manifold and their distances on the manifold are computed [2]. In this work, $k = 2$.

$$M_{i,j,k,5} = Q_{E(G)}\left(\sum_{\kappa \in \backslash knn(k)} \left|\widehat{Y_{\kappa,(\iota,\jmath),w}} - Y_{\kappa}\right|\right)$$

Supplementary Equation 5. Formula for the local error metafeature.

### Theoretical Models

We simulated the following data sets, each with sample size $n = 60$, entity measurement set 1 size $|S_1| = 11$, and entity measurement set 2 size $|S_2| = 6$:

- **Single Continuous:** 30 data sets with continuous outcome in which a single feature association *(a,b)* provided the optimal predictor. Data generation is described in Supplementary Equation 6, where $\mathfrak{N}$ is the noise level, $X^{S_1}$ is the matrix of measurements for entity set 1, $X^2$ is the matrix of measurements for entity set 2, $Y$ is the outcome vector, $\epsilon$ is the noise variable, $\sigma(X)$ is the standard deviation of $X$, and $N(0,1)$ is the normal distribution. Noise levels of 0, 1, 0.1, and 0.5 were simulated.

$$X^{S_1}, \tilde{X}_j^{S_2}, Y \sim N(0,1), j \neq b$$

$$\tilde{X}_b^{S_2} = \frac{X_a^{S_1}}{Y}$$

$$X_j^{S_2} = \tilde{X}_j^{S_2}, j \neq b$$
$$\epsilon = N(0, \mathfrak{N} \times \sigma(\tilde{X}^{S_2}))$$
$$X_b^{S_2} = \tilde{X}_b^{S_2} + \epsilon$$

Supplementary Equation 6. Data generation for the Single Continuous case.

- **Single Discrete:** 30 data sets with discrete outcome in which a single feature association *(a,b)* provided the optimal predictor. Data generation is described in Supplementary Equation 7, where $\mathfrak{B}(0.5)$ is the binomial distribution where all outcomes have equal probability. Noise levels of 0, 0.1, and 0.5 were simulated.

$$X^{S_1}, \tilde{X}_j^{S_2}, Y \sim N(0,1), j \neq b$$
$$\mathrm{Y} = \mathfrak{B}(0.5)$$
$$\tilde{X}_b^{S_2} = \frac{X_a^{S_1}}{Y}$$
$$X_j^{S_2} = \tilde{X}_j^{S_2}, j \neq b$$
$$\epsilon = N(0, \mathfrak{N} \times \sigma(\tilde{X}^{S_2}))$$
$$X_b^{S_2} = \tilde{X}_b^{S_2} + \epsilon$$

Supplementary Equation 7. Data generation for the Single Discrete case.

- **Complementary Continuous:** 30 data sets with continuous outcome in which a pair of feature associations *(a, b)* and *(c, d)* combine to produce an optimal predictor. Data generation is described in Supplementary Equation 8. Noise levels of 0.01, 0.1, and 0.5 were simulated.

$$X^{S_1}, X_j^{S_2}, Y \sim N(0,1), j \notin \{b, d\}$$
$$\tilde{X}_b^{S_2} = \frac{X_a^{S_1}}{Y}, \tilde{X}_d^{S_2} = \frac{X_c^{S_1}}{Y}$$
$$X_j^{S_2} = \tilde{X}_j^{S_2}, j \notin \{b, d\}$$
$$\epsilon = N(0, \mathfrak{N} \times \sigma(\tilde{X}^{S_2}))$$
$$X_b^{S_2} = \tilde{X}_b^{S_2} + \epsilon, X_d^{S_2} = \tilde{X}_d^{S_2} - \epsilon$$

Supplementary Equation 8. Data generation for the Complementary Continuous case.

- **Complementary Discrete:** 30 data sets with discrete outcome in which a pair of feature associations *(a, b)* and *(c, d)* combine to produce an optimal predictor. Data generation is described in Supplementary Equation 9. Noise levels of 0.01, 0.1, and 0.5 were simulated.

$$X^{S_1}, \tilde{X}_j^{S_2}, Y \sim N(0,1), j \notin \{b, d\}$$
$$\mathrm{Y} = \mathfrak{B}(0.5)$$
$$\tilde{X}_b^{S_2} = \frac{X_a^{S_1}}{Y}, \tilde{X}_d^{S_2} = \frac{X_c^{S_1}}{Y}$$
$$X_j^{S_2} = \tilde{X}_j^{S_2}, j \notin \{b, d\}$$
$$\epsilon = N(0, \mathfrak{N} \times \sigma(\tilde{X}^{S_2}))$$
$$X_b^{S_2} = \tilde{X}_b^{S_2} + \epsilon, X_d^{S_2} = \tilde{X}_d^{S_2} - \epsilon$$

Supplementary Equation 9. Data generation for the Single Continuous case.

### Ablation Models

GNN + CMP is identical to GENN, but without SigProp edge pruning, i.e., $G_{opt} = G$, so that only ϕ is learned. Evaluating GENN against GNN + CMP highlights the effect of SigProp pruning on model performance. The training process for GNN + CMP is described in Supplementary Algorithm 3.

Further, GNN + SigProp is identical to GENN, but the predictions of each component predictor are combined using averaging, mimicking a standard graph pooling procedure or weighted ensemble model (Supplementary Equation 10) [3]. Evaluating GENN against GNN + SigProp highlights the effect of Composite Model Pooling.

$$\widehat{Y_{k,G,w}} = \sum_{(i,j)\in E(G)} w_{(i,j),k} \times \frac{\left(X_{i,k}^{S_1} - \left(\beta_{0,i,j} + X_{j,k}^{S_2}\beta_{1,i,j} + \sum_{c=1}^{|S_3|}\left(X_{c,k}^{S_3}\beta_{3+c,i,j}\right)\right)\right)}{\left(\beta_{2,i,j} + X_{j,k}^{S_2}\beta_{3,i,j}\right)}$$

Supplementary Equation 10. GNN + SigProp composite predictor.

The gradient of ϕ with respect to GNN + SigProp is given in Supplementary Equation 11.

$$\nabla_{\phi} = \frac{\partial(\epsilon)}{\partial(\phi)} = \sum_{k=1}^{n} (2(\widehat{Y_{k,G_{opt},\phi}} - Y)(M_{i,j,k} \times \sum_{(i,j)\in E(G)} \frac{Num}{Denom})$$

$$Num = X_{i,k}^{S_1} - \left(\beta_{0,i,j} + X_{j,k}^{S_2}\beta_{1,i,j} + \sum_{c=1}^{|S_3|}\left(X_{c,k}^{S_3}\beta_{3+c,i,j}\right)\right)$$

$$Denom = \left(\beta_{2,i,j} + X_{j,k}^{S_2}\beta_{3,i,j}\right)$$

Supplementary Equation 11. Gradient for training GNN + SigProp.

### Hyperparameters

For the theoretical models, GENN models were trained with learning rate $\eta = 0.5$ with no $R^2$ filtering. Random Forests were trained with 500 trees, and *mtry* (number of entities available to Random Forest during each training step) tuned using 10-fold cross-validation ranging from $\{1, \ldots, |S1| + |S2|\}$ by $\frac{(|S1|+|S2|)}{10}$. LASSO and Group LASSO $\lambda$ values (weight of the LASSO penalty term) were also tuned using 10-fold cross-validation ranging from $\{0.01, \ldots, 0.20\}$ by 0.01, and the groups in Group LASSO were the 1-hop neighborhoods of the graph.

For the NCI-60 and TOPMed data sets, $\eta$ between $\{0.1, \ldots, 0.9\}$ were tested in increments of 0.1 on only a small subgraph of the full graph filtered using stringent $R^2$ cutoffs of 0.7 and 0.1, respectively (these were chosen visually; see **Supplementary Figures 2-3**). Results can be found in **Supplementary Table 18**. After selecting $\eta$, $R^2$ cutoffs between $\{0.3, \ldots, 0.7\}$ were tested in increments of 0.1 on the NCI-60 data set $R^2$ cutoffs between $\{0.01, \ldots, 0.1\}$ were tested in increments of 0.01 on the TOPMed data set. Results shown in the manuscript for are for the best-performing $R^2$, $\lambda$, or *mtry* cutoffs for the NCI-60 and TOPMed data **(Supplementary Tables 19-21).** For the TCGA data, we set $\eta = 0.2$, $R^2 > 0.2$, and FDR-adjusted $p < 0.2$.

### Advantage of Scaled Covariance Metric

$SCov$ accounts for predictor range in a manner other regression metrics do not, as illustrated using the following models: (1) A model that always predicts the exact mean of the data, i.e., $\widehat{Y_1} = \bar{Y}$, (2) A model where the predicted values increase or decrease with the true values, but on a different scale, i.e., $\widehat{Y_2} = \bar{Y} + \varepsilon(Y - \bar{Y})$, and (3) A model that predicts exactly the correct value, i.e., $\widehat{Y_4} = Y$. Model 1 is equivalent to model 2 when $\varepsilon = 0$, and model 3 is equivalent to model 2 when $\varepsilon = 1$. When $0 < \varepsilon < 1$ or $1 < \varepsilon$, the trend of $\widehat{Y_2}$ mirrors the trend of $Y_2$, but the ranges differ by the scale $\varepsilon$. A metric $m(Y, \hat{Y})$ where higher is "better" should therefore reflect the criterion given in Supplementary Equation 13.

$$m(Y, \widehat{Y_1}) < (m(Y, \widehat{Y_2}), \varepsilon > 0, \varepsilon \neq 1) < m(Y, \widehat{Y_3})$$

Supplementary Equation 13. Mathematical criterion for measuring the performance of a model.

Absolute Error based metrics, Squared Error based metrics, correlation, and covariance do not meet this criterion, but $SCov$ does, as illustrated below.

**Sum Absolute Error** is defined as $\sum_{k=1}^{n} |\hat{Y}[k] - Y[k]|$.
Therefore, $m(Y, \widehat{Y_1}) = 1 - \sum_{k=1}^{n} |Y[k] - \bar{Y}|$.
Further, $m(Y, \widehat{Y_2}, \varepsilon = 2) = 1 - \sum_{k=1}^{n} |Y[k] - (\bar{Y} + 2Y[k] - 2\bar{Y})|$.
Simplification yields $m(Y, \widehat{Y_2}, \varepsilon = 2) = 1 - \sum_{k=1}^{n} |\bar{Y} - Y[k]| = 1 - \sum_{k=1}^{n} |Y[k] - \bar{Y}|$.
Therefore, $m(Y, \widehat{Y_2}) = m(Y, \widehat{Y_1})$, violating the criterion.

**Sum Squared Error** is defined as $\sum_{k=1}^{n} (\hat{Y}[k] - Y[k])^2$.
Therefore, $m(Y, \widehat{Y_1}) = 1 - \sum_{k=1}^{n} (Y[k] - \bar{Y})^2$.
Further, $m(Y, \widehat{Y_2}, \varepsilon = 2) = 1 - \sum_{k=1}^{n} \left(Y[k] - (\bar{Y} + 2Y[k] - 2\bar{Y})\right)^2$.
Simplification yields $m(Y, \widehat{Y_2}, \varepsilon = 2) = 1 - \sum_{k=1}^{n} (\bar{Y} - Y[k])^2 = 1 - \sum_{k=1}^{n} (Y[k] - \bar{Y})^2$.
Therefore, $m(Y, \widehat{Y_2}) = m(Y, \widehat{Y_1})$, violating the criterion.

**Covariance** $Cov$ is defined as $\sum_{k=1}^{n} \frac{(\hat{Y}[k] - \bar{\hat{Y}})(Y[k] - \bar{Y})}{n}$.

Therefore, $m(Y, \widehat{Y_2}) = \sum_{k=1}^{n} \frac{\left(\left(\bar{Y} + \varepsilon(Y[k] - \bar{Y}) - \sum_{l=1}^{n} \frac{\bar{Y} + \varepsilon(Y[k] - \bar{Y})}{n}\right\}\right)(Y[k] - \bar{Y})}{n}$.

Distribution yields $m(Y, \widehat{Y_2}) = \sum_{k=1}^{n} \frac{\left(\bar{Y} + \varepsilon(Y[k] - \bar{Y}) - \sum_{l=1}^{n} \frac{\bar{Y}}{n} + \varepsilon \sum_{l=1}^{n} \frac{Y[l]}{n} - \varepsilon \sum_{l=1}^{n} \frac{\bar{Y}}{n}\right)(Y[k] - \bar{Y})}{n}$.

Simplification yields $m(Y, \widehat{Y_2}) = \sum_{k=1}^{n} \frac{(\bar{Y} + \varepsilon(Y[k] - \bar{Y}) - \bar{Y})(Y[k] - \bar{Y})}{n} = \sum_{k=1}^{n} \frac{\varepsilon(Y[k] - \bar{Y})(Y[k] - \bar{Y})}{n}$.

Now, $m(Y, \widehat{Y_3}) = \sum_{k=1}^{n} \frac{(Y[k] - \bar{Y})(Y[k] - \bar{Y})}{n}$.
So, $m(Y, \widehat{Y_2}) = \varepsilon \times m(Y, \widehat{Y_3})$.
Then, if $\varepsilon > 1$, $m(Y, \widehat{Y_2}) > m(Y, \widehat{Y_3})$, violating the criterion.

**Correlation** is defined as $\frac{Cov(Y, \hat{Y})}{\sum_{k=1}^{n} (Y[k] - \bar{Y}) \times \sum_{k=1}^{n} (\hat{Y}[k] - \bar{\hat{Y}})}$.

$Cov(Y, \widehat{Y_2}) = \varepsilon \times Cov(Y, \widehat{Y_3})$.
Now, it was also previously shown that $\sum_{k=1}^{n} \left(\hat{Y}[k] - \bar{\hat{Y}}\right) = \varepsilon \sum_{k=1}^{n} (Y[k] - \bar{Y})$.
Therefore, $\frac{Cov(Y, \widehat{Y_2})}{\sum_{k=1}^{n} (Y[k] - \bar{Y}) \times \sum_{k=1}^{n} (\widehat{Y_2}[k] - \overline{\widehat{Y_2}})} = \frac{\varepsilon \times Cov(Y, \widehat{Y_3})}{\varepsilon \sum_{k=1}^{n} (Y[k] - \bar{Y}) \times \sum_{k=1}^{n} (Y[k] - \bar{Y})}$

Further, $\sum_{k=1}^{n}\left(\hat{Y}[k] - \bar{\hat{Y}}\right) = \sum_{k=1}^{n}(Y[k] - \bar{Y})$.
So $m\left(Y, \widehat{Y_2}\right) = m\left(Y, \widehat{Y_3}\right)$, violating the criterion.
SCov is defined as $\frac{Cov(Y,\hat{Y})}{\max\{(\sum_{k=1}^{n}(Y[k]-\bar{Y}),\sum_{k=1}^{n}(\hat{Y}[k]-\bar{\hat{Y}})\})}$, or the covariance scaled by the square of the maximal variance between the true outcome and the predicted outcome. Scaling by the maximal variance allows differences in true and predicted outcome range to be factored into the performance metric.
$Cov\left(m\left(Y, \widehat{Y_1}\right)\right) = 0$, so $SCov\left(m\left(Y, \widehat{Y_1}\right)\right) = 0$.
Furthermore, for $\varepsilon < 1$, $\sum_{k=1}^{n}\left(\hat{Y}[k] - \bar{\hat{Y}}\right) < \sum_{k=1}^{n}(Y[k] - \bar{Y})$, so $SCov\left(m\left(Y, \widehat{Y_2}\right)\right) = \frac{\varepsilon \times Cov(Y,\hat{Y})}{\left(\sum_{k=1}^{n}(Y[k]-\bar{Y})\right)^2} < \frac{Cov(Y,\hat{Y})}{\left(\sum_{k=1}^{n}(Y[k]-\bar{Y})\right)^2} = SCov\left(m\left(Y, \widehat{Y_3}\right)\right)$.
Furthermore, for $\varepsilon > 1$, $\sum_{k=1}^{n}\left(\hat{Y}[k] - \bar{\hat{Y}}\right) > \sum_{k=1}^{n}(Y[k] - \bar{Y})$, so $SCov\left(m\left(Y, \widehat{Y_2}\right)\right) = \frac{\varepsilon \backslash Cov(Y,\hat{Y})}{\left(\varepsilon \sum_{k=1}^{n}(Y[k]-\bar{Y})\right)^2} < \frac{\backslash Cov(Y,\hat{Y})}{\left(\sum_{k=1}^{n}(Y[k]-\bar{Y})\right)^2} = SCov\left(m\left(Y, \widehat{Y_3}\right)\right)$.
Additionally, because $\left|Y[k] - \widehat{Y_2}[k]\right| < |Y[k] - \bar{Y}[k]|$ for all $k \in Set(1 \ldots n)$, $Cov\left(Y, \widehat{Y_2}\right) > 0$.
Therefore, $SCov\left(Y, \widehat{Y_1}\right) < SCov\left(Y, \widehat{Y_2}\right) < SCov\left(Y, \widehat{Y_3}\right)$.

### Constructing Literature-Supported Analyte Groups for NCI-60

To construct the set of analytes and analyte groups supported by literature $V_{lit}$ with respect to the NCI-60 pan-cancer database, literature was retrieved using the following queries in PubMed. $V_{lit}$ was compiled as described below using [4-6].

- "5 fluorouracil mechanism"
- "metabolomics + 5-fu + pan-cancer"
- "pathways + 5-fu – gastric – colorectal – colon – pancreatic – ovarian – gastrointestinal – carcinoma – cervical – esophageal – epithelial".

To match metabolites which do not have standardized names, the metabolite names used in $V_{lit}$ were manually mapped to the metabolite names.
To find $V_{opt} \cap V_{lit}$, R was used to match the gene and metabolite names to those from the literature. To match pathways, the *bitr*() function from the *clusterProfiler* R package and the *runCombinedFisherTest*() function from the *RaMP-DB* R package, version 2.3.0, were used. To match chemical classes, the *chemicalClassEnrichment*() function from the *RaMP-DB* R package was used. Pathway and metabolite class names were manually mapped from the R package function results to $V_{lit}$.


1. Allen, M.P., *The t test for the simple regression coefficient*, in *Understanding Regression Analysis*. 1997, Springer Science + Business. p. 66-70.
2. Ding, H., et al., *Integrative cancer patient stratification via subspace merging.* Bioinformatics, 2019. **35**: p. 1653-1659.
3. Dietterich, T.G., *Ensemble Methods in Machine Learning*. Vol. 1857. 2000. 1-15.
4. Braisted, J., et al., *RaMP-DB 2.0: a renovated knowledgebase for deriving biological and chemical insight from metabolites, proteins, and genes.* Bioinformatics, 2023. **39**(1).

5. Yu, G., et al., *clusterProfiler: an R package for comparing biological themes among gene clusters.* OMICS, 2012. **16**(5): p. 284-7.
6. Carlson, M., *org.Hs.eg.db: Genome wide annotation for Human*. 2019.
