## Supplementary References for "Finding Salient Multi-Omic Interactomes Relevant to Multiple Biomedical Outcomes using Graph Ensemble Neural Networks"

### 1. Methods that link analytes associations or groups to outcome *a posteriori* or model outcome-dependent associations:

Commented [Ma1]: AB: I suggest adding numbers and calling them on the main text. I think it makes it more friendly.


Xiang, S., et al., *Mining of differentially co-expressed gene modules reveals perturbed functional network and key regulators in Alzheimer's disease brains.* Alzheimer's & Dementia, 2020. **16**: p. e047095.

Ideker, T. and N.J. Krogan, *Differential network biology.* Molecular Systems Biology, 2012. **8**: p. 565.

Siddiqui, J.K., et al., *IntLIM: Integration using linear models of metabolomics and gene expression data.* BMC Bioinformatics, 2018. **19**: p. 81.

Eicher, T., et al., *IntLIM 2.0: identifying multi-omic relationships dependent on discrete or continuous phenotypic measurements.* Bioinformatics Advances, 2023. **3**(1): p. vbad009.

Siska, C., R. Bowler, and K. Kechris, *The discordant method: a novel approach for differential correlation.* Bioinformatics, 2016. **32**: p. 690-6.

Fukushima, A., *DiffCorr: An R package to analyze and visualize differential correlations in biological networks.* Gene, 2013. **518**: p. 209-214.

Shi, W.J., et al., *Unsupervised discovery of phenotype-specific multi-omics networks.* Bioinformatics, 2019. **35**: p. 4336-4343.

Ma, J., et al., *Differential network enrichment analysis reveals novel lipid pathways in chronic kidney disease.* Bioinformatics, 2019. **35**(18): p. 3441–3452.

Langfelder, P. and S. Horvath, *WGCNA: An R package for weighted correlation network analysis.* BMC Bioinformatics, 2008. **9**.

Smyth, G.K., *Linear models and empirical bayes methods for assessing differential expression in microarray experiments.* Statistical Applications in Genetics and Molecular Biology, 2004. **3**.

Shi, W.J., et al., *Unsupervised discovery of phenotype-specific multi-omics networks.* Bioinformatics, 2019. **35**: p. 4336-4343.

Fukushima, A., *DiffCorr: An R package to analyze and visualize differential correlations in biological networks.* Gene, 2013. **518**: p. 209-214.

Siska, C., R. Bowler, and K. Kechris, *The discordant method: a novel approach for differential correlation.* Bioinformatics, 2016. **32**: p. 690-6.

Ma, J., et al., *Differential network enrichment analysis reveals novel lipid pathways in chronic kidney disease.* Bioinformatics, 2019. **35**(18): p. 3441–3452.


### 2. Methods for determining node importance in Random Forests:


Liaw, A., *Breiman and Cutler's Random Forests for Classification and Regression*. 2018, The Comprehensive R Archive Network.

Liaw, A. and M. Wiener, *Classification and Regression by randomForest*, in *R News*. 2002.

Archer, K.J. and R.V. Kimes, *Empirical characterization of random forest variable importance measures.* Computational Statistics and Data Analysis, 2008. **52**: p. 2249-2260.

Breiman, L., *Random forests.* Machine Learning, 2001. **45**: p. 5-32.

Kursa, M. B., A. Jankowski and W. R. Rudnicki (2010). "Boruta – A System for Feature Selection." Fundamenta Informaticae **101**(4): 271-285.

### 3. Methods for learning large Bayesian Networks:

Scanagatta, M., et al., *Learning Bayesian Networks with Thousands of Variables*, in *Twenty-Ninth Conference on Neural Information Processing Systems*. 2015.

Cussens, J., et al. *Bayesian Network Structure Learning with Integer Programming: Polytopes, Facets and Complexity (Extended Abstract)*. in *International Joint Conference on Artificial Intelligence*. 2017.

Teyssier, M. and D. Koller. *Ordering-based search: a simple and effective algorithm for learning Bayesian networks*. in *Uncertainty in Artificial Intelligence*. 2005.

### 4. Graph Neural Network methods for graph classification:

Zhang, M., et al. *An End-to-End Deep Learning Architecture for Graph Classification*. in *AAAI Conference on Artificial Intelligence*. 2018.

Kong, Y. and T. Yu, *forgeNet: a graph deep neural network model using tree-based ensemble classifiers for feature graph construction.* Bioinformatics, 2020. **36**: p. 3507-3515.

Lee, J.B., R. Rossi, and X. Kong, *Graph Classification using Structural Attention*. 2018.

Kong, Y. and T. Yu, *A graph-embedded deep feedforward network for disease outcome classification and feature selection using gene expression data.* Bioinformatics, 2018. **34**(21): p. 3727-3737.

### 5. Methods to identify the salient subgraph in Graph Neural Networks:

Baldassarre, F. and H. Azizpour, *Explainability Techniques for Graph Convolutional Networks.* ArXiv, 2019.

Sanchez-Lengeling, B., et al., *Evaluating Attribution for Graph Neural Networks*, in *Neural Information Processing Systems*. 2020. p. 58-62.

Smilkov, D., et al., *SmoothGrad: Removing Noise by Adding Noise.* ArXiv, 2017.

Schwarzenberg, R., et al., *Layerwise Relevance Visualization in Convolutional Text Graph Classifiers*, in *Graph-Based Methods for Natural Language Processing*. 2019.

Ying, R., et al., *GNNExplainer: Generating Explanations for Graph Neural Networks*, in *Neural Information Processing Systems*. 2019.

Luo, D., et al., *Parameterized Explainer for Graph Neural Network.* ArXiv, 2020.

Schlichtkrull, M.S., N. De Cao, and I. Titov, *Interpreting Graph Neural Networks for NLP with Differentiable Edge Masking*, in *International Conference on Learning Representations*. 2021.

Wang, X., et al., *Reinforced Causal Explainer for Graph Neural Networks.* IEEE Transactions on Pattern Analysis and Machine Intelligence, 2022. **45**(2).

Lucic, A., et al., *CF-GNNExplainer: Counterfactual Explanations for Graph Neural Networks*, in *International Conference on Artificial Intelligence and Statistics*. 2022.

### 6. Methods for graph segmentation:


Bianchi, F.M., D. Grattarola, and C. Alippi. *Spectral Clustering with Graph Neural Networks for Graph Pooling*. in *37th International Conference on Machine Learning*. 2019. International Machine Learning Society (IMLS).

Gao, H. and S. Ji, *Graph U-Nets.* IEEE Transactions on Pattern Analysis and Machine Intelligence, 2022. **44**: p. 4948-4960.

Zhang, M., et al., *An End-to-End Deep Learning Architecture for Graph Classification.* 2018: p. 4438-4445.

Ying, Z., et al., *Hierarchical Graph Representation Learning with Differentiable Pooling*, in *Advances in Neural Information Processing Systems 31*. 2018.

Lee, J., I. Lee, and J. Kang. *Self-Attention Graph Pooling*. in *36th International Conference on Machine Learning*. 2019.


### 7. Other Graph Neural Network references:


Veličković, P., et al., *Graph Attention Networks*, in *6th International Conference on Learning Representations*. 2018, International Conference on Learning Representations, ICLR.

Liu, C., et al., *Graph Pooling for Graph Neural Networks: Progress, Challenges, and Opportunities.* ArXiv, 2023.

Niepert, M., M. Ahmed, and K. Kutzkov, *Learning Convolutional Neural Networks for Graphs*, in *33rd International Conference on Machine Learning*. 2016.

Yang, Y. and D. Li. *NENN: Incorporate Node and Edge Features in Graph Neural Networks*. in *12th Asian Conference on Machine Learning*. 2020.

Wang, Z., J. Chen, and H. Chen, *EGAT: Edge-Featured Graph Attention Network*, in *30th International Conference on Artificial Neural Networks*. 2021. p. 253-264.

Gong, L. and Q. Cheng, *Exploiting Edge Features for Graph Neural Networks*, in *IEEE / CVF Computer Vision and Pattern Recognition Conference*. 2019. p. 9211-9219.


### 8. Corpora of literature specific to 5-FU uptake mechanisms:


Ghafouri-Fard, S., et al., *5-Fluorouracil: A Narrative Review on the Role of Regulatory Mechanisms in Driving Resistance to This Chemotherapeutic Agent.* Frontiers in Oncology, 2021. **11**: p. 658636.

Danesh Pouya, F., Y. Rasmi, and M. Nemati, *Signaling Pathways Involved in 5-FU Drug Resistance in Cancer.* Cancer Investigation, 2022. **40**(6): p. 516-543.

Miura, K., et al., *5-FU Metabolism in Cancer and Orally-Administrable 5-FU Drugs.* Cancers, 2010. **2**(3): p. 1717-1730.

Scartozzi, M., et al., *5-fluorouracil pharmacogenomics: still rocking after all these years?* Pharmacogenomics, 2011. **12**(2): p. 251-265.

Kang, J., et al., *Ribosomal proteins and human diseases: molecular mechanisms and targeted therapy.* Signal Transduction and Targeted Therapy, 2021. **6**(1): p. 323.

Lam, S.W., H.J. Guchelaar, and E. Boven, *The role of pharmacogenetics in capecitabine efficacy and toxicity.* Cancer Treatment Reviews, 2016. **50**: p. 9-22.

Rodrigues, D., et al., *New insights into the mechanisms underlying 5-fluorouracil-induced intestinal toxicity based on transcriptomic and metabolomic responses in human intestinal organoids.* Archives of Toxicology, 2021. **95**(8): p. 2691-2718.

Sanchez-Castillo, A., M. Vooijs, and K.R. Kampen, *Linking Serine/Glycine Metabolism to Radiotherapy Resistance.* Cancers (Basel), 2021. **13**(6).

Costa-Pinheiro, P., et al., *Diagnostic and prognostic epigenetic biomarkers in cancer.* Epigenomics, 2015. **7**(6): p. 1003-1015.

Cao, D. and G. Pizzorno, *Uridine phosophorylase: an important enzyme in pyrimidine metabolism and fluoropyrimidine activation.* Drugs Today (Barc), 2004. **40**(5): p. 431-43.
