## Supplementary Figures for "Finding Salient Multi-Omic Interactomes Relevant to Multiple Biomedical Outcomes using Graph Ensemble Neural Networks"

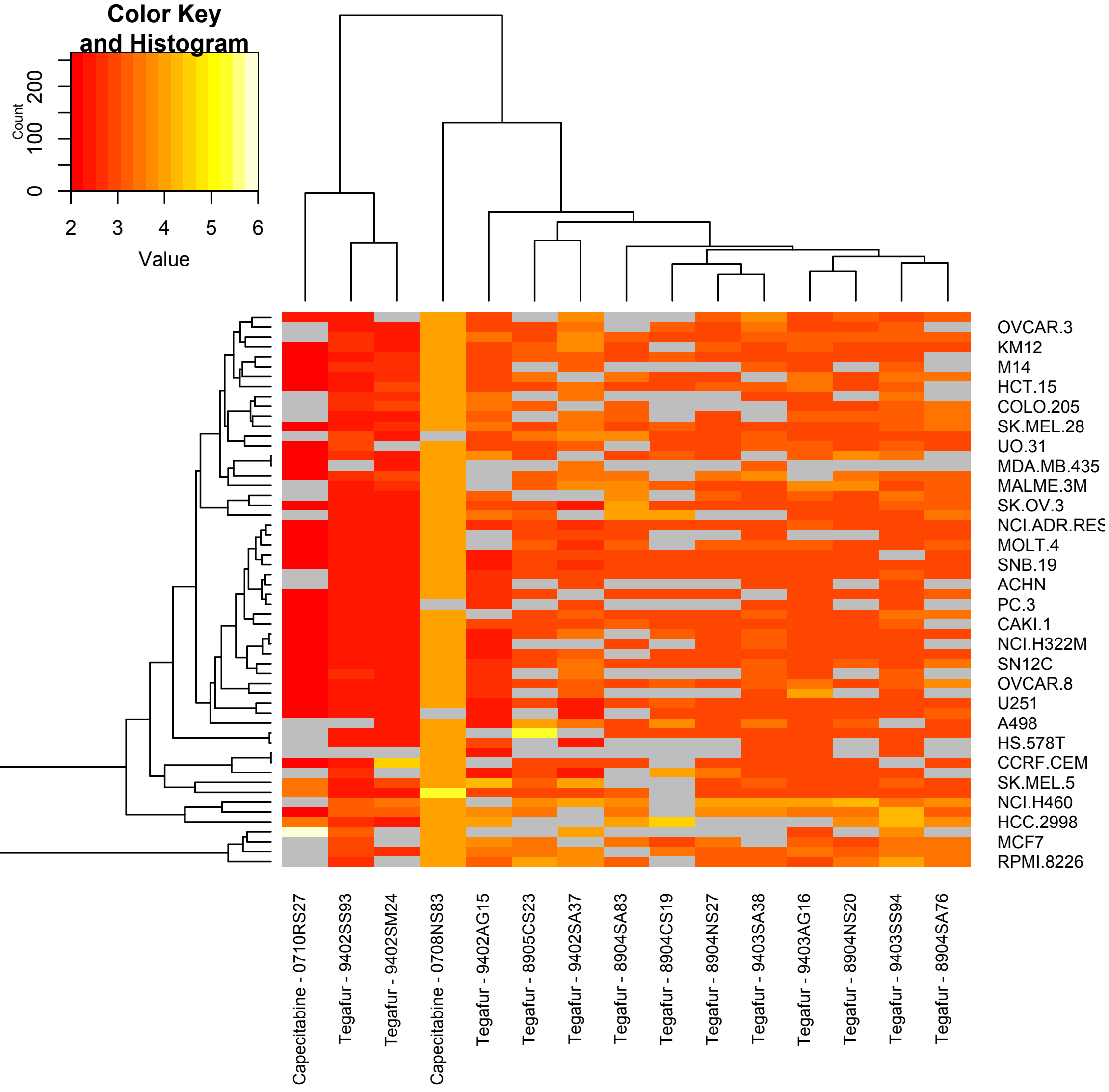


Supplementary Figure 1. Several of the 5-FU drug response experiments exhibited little to no $IC_{50}$ variability across cell lines.

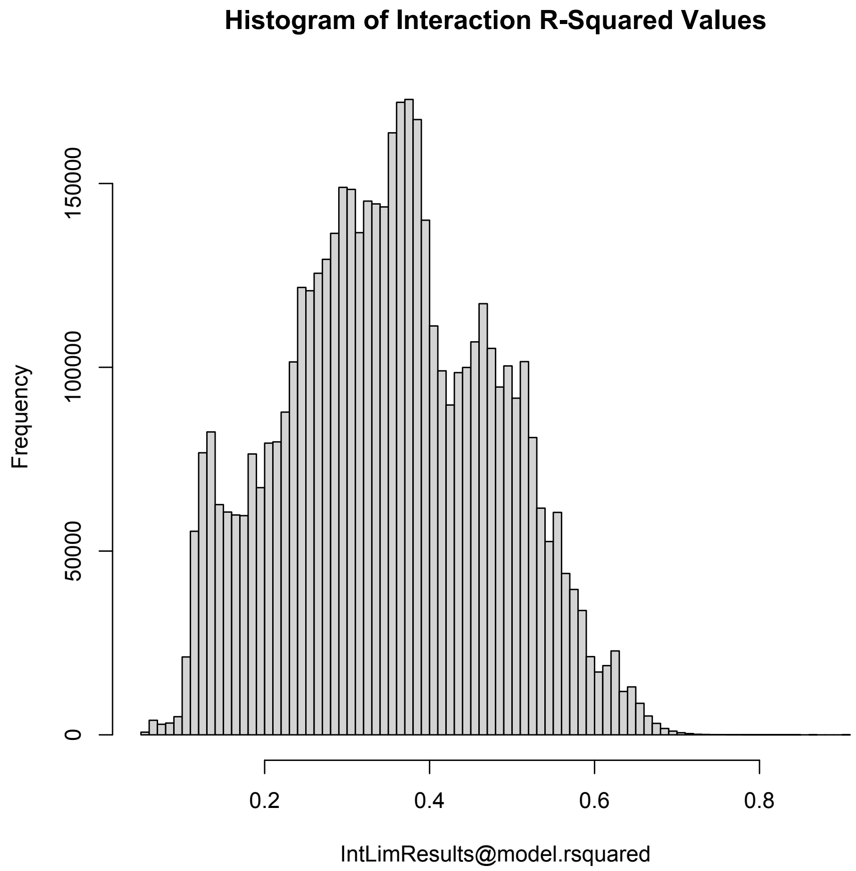


Supplementary Figure 2. $R^2$ distribution across all regression models from the NCI-60 data set.

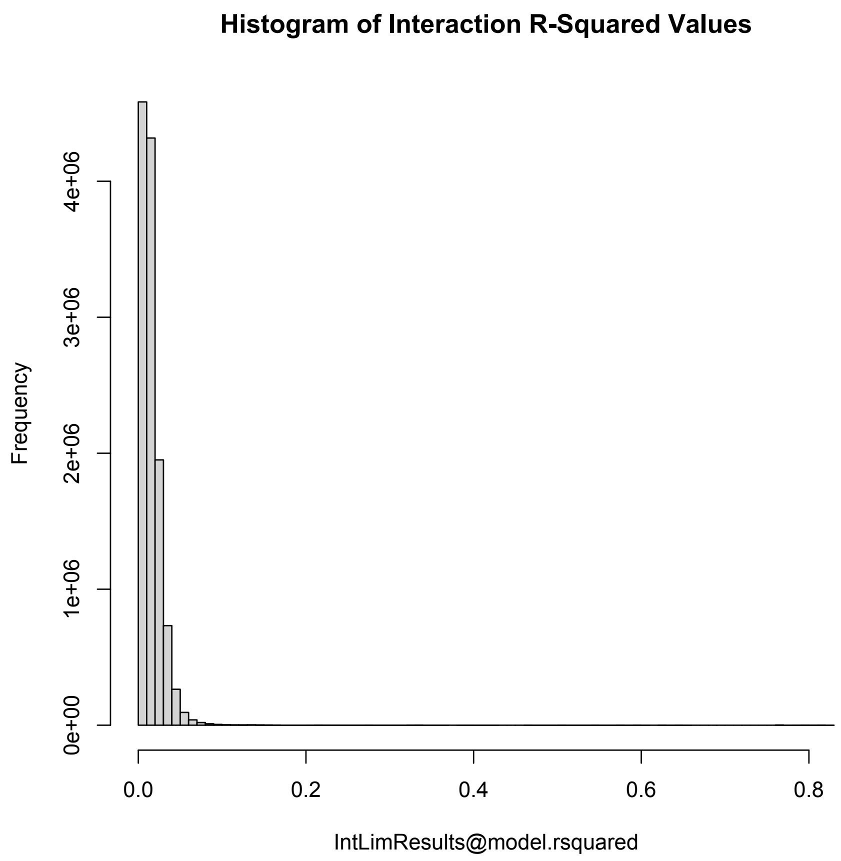


Supplementary Figure 3. $R^2$ distribution across all regression models from the GACRS data set.
